# Multi-Trait Meta-QTL Analysis Reveals Genomic Hotspot Classes for Strategic Maize Improvement

**DOI:** 10.64898/2026.06.06.730627

**Authors:** Seshasai Parthasarathy, Klaus Koehler, Torbert Rocheford

## Abstract

**Background:** Decades of maize (*Zea mays* L.) quantitative trait locus (QTL) mapping have produced fragmented results across hundreds of independent studies, characterized by broad confidence intervals, population-specific effects, and a predominantly single-trait scope. Comprehensive multi-trait integration remains limited in the public domain, yet it could improve our understanding of trait relationships for strategic maize breeding. We integrated 2,701 QTLs published over 30 years across five trait categories (grain yield and components (GYC); plant development and architecture (PDA); plant physiology and stress adaptation (PHS); grain quality and nutritional composition (GQN); and disease and pest resistance (DPR)) to identify functionally classified genomic hotspots and prioritize candidate genes for multi-trait breeding.

**Results:** BioMercator V4.2.3 consolidated 2,518 projectable QTLs into 187 high-confidence meta-QTLs (MQTLs), achieving a 4.0-fold (75%) reduction in mean genetic-map confidence interval width (47.6 to 11.8 cM); 128 of 187 MQTLs (68.4%) showed genome-wide association study (GWAS) co-localization, and 119 (63.6%) met the strict dual-platform criterion. Twenty-three genomic hotspots harbored 131 of 187 MQTLs (70.1%) and were classified by an ordered, data-driven rule into three categories: ten multi-trait hubs enabling simultaneous improvement through pleiotropic or tightly linked genes; seven pathway-focused clusters dominated by a single trait pathway, exemplified by the chromosome 8 kernel-yield cluster; and six major-effect loci, each carrying a member meta-QTL of large effect (phenotypic variance explained (PVE) of at least 28%), including the Rp1 common-rust locus (35% member-MQTL PVE) and the chromosome 9 Wx1/THP9 starch-and-protein locus (up to 52% member-MQTL PVE). Environmental classification distinguished MQTLs predominantly supported by optimal-condition QTLs (42%) from those supported by stress-condition QTLs (28%), the latter showing ∼1.7-fold greater mean contributing-QTL phenotypic variance (12.8% vs. 7.4%), consistent with conditional effect amplification under stress; within-population comparisons under contrasting water regimes showed a larger, ∼3.5-fold amplification. Network-based candidate gene prioritization with cross-cereal ortholog analysis identified priority candidates with confirmed orthologs in one or more comparison cereals (rice, sorghum, wheat, or barley), with conservation highest for developmental and metabolic genes and lowest for pathogen-resistance genes, identifying priority targets for functional genomics investment.

**Conclusions:** This functionally classified and environmentally characterized meta-QTL framework provides breeders with a structured resource for multi-trait hotspot selection, environment-appropriate allele deployment, and functional genomics prioritization, with broader applicability as a transferable template for other crops confronting fragmented QTL literature and complex multi-trait breeding objectives.

## Background

Maize (*Zea mays* L.) is one of the world’s most important cereal crops. It carries extensive genetic diversity that reflects complex genetic control of many traits and that was shaped by domestication from wild teosinte followed by intensive modern breeding across diverse environments [1–2]. This diversity provides broad opportunities for crop improvement; however, translating accumulated genetic knowledge into tangible breeding progress requires a clearer understanding of how the genomic regions controlling key agronomic traits are organized and distributed across diverse genetic backgrounds and environmental conditions.

Over the past three decades, traditional linkage mapping in biparental populations has identified thousands of quantitative trait loci (QTLs) affecting important maize traits [3], and genome-wide association studies (GWAS) using diverse germplasm panels have subsequently provided enhanced mapping resolution and broader genetic coverage across the species’ diversity [4–6]. Despite this wealth of genetic information, several challenges limit the efficient practical application of mapping results in breeding programs. Many QTLs are detected only in specific genetic backgrounds or environmental conditions, whereas others span broad genomic intervals containing hundreds of candidate genes that are difficult to prioritize without additional functional information [3–4]. Initial QTL effect-size estimates can be inflated owing to sampling variation and publication bias, a pattern formalized as the Beavis effect [7–8], whereby QTLs reaching statistical significance in small mapping populations systematically overestimate true effect sizes, creating expectations that are often not replicated when loci are deployed in breeding. Extensive genotype-by-environment interactions further complicate the identification of broadly relevant loci versus population– or environment-specific artifacts [9]. Consequently, despite decades of intensive mapping research, relatively few QTLs have been fine-mapped to causal genes or effectively deployed in operational plant breeding programs [10–12].

Meta-QTL analysis was designed to address these challenges by synthesizing QTL information across multiple independent studies, distinguishing consistent genetic signals from study-specific variation, and substantially reducing confidence intervals through precision-weighted averaging across independent populations and genetic backgrounds [13–14]. This approach has demonstrated meaningful confidence-interval reductions across diverse crop species while revealing patterns of genetic architecture not apparent from individual studies conducted in isolation [15–19]. In maize specifically, meta-analyses have provided valuable insights for drought tolerance [20–21], disease resistance [22–23], yield components [24], grain quality [25–26], and physiological traits [27]. However, previous maize meta-analyses carry noteworthy limitations that restrict their practical utility for breeding programs targeting multiple traits simultaneously. Their predominantly single-trait scope omits critical information on co-localizations among traits that is needed for understanding genetic correlations, anticipating unintended selection consequences across traits, and identifying loci capable of simultaneously improving multiple breeding objectives [28–29]. A genomic region influencing drought tolerance may, for example, also substantially affect yield potential under optimal conditions, creating trade-offs that single-trait analysis cannot detect or quantify [30–31]. Without multi-trait integration, breeders cannot easily distinguish loci suitable for independent manipulation from those requiring strategic multi-objective selection decisions informed by the pleiotropy or tight linkage governing co-localizing effects on different traits [28, 32–34].

Most existing meta-analyses have also lacked systematic verification through association mapping, despite the well-established complementary strengths of linkage and association approaches [35–38]. Linkage mapping in biparental populations provides high statistical power but typically yields broad confidence intervals (commonly 20–40 Mb), owing to the limited recombination captured in small populations [4–5]. GWAS in diversity panels, by contrast, achieves finer mapping resolution by exploiting the historical recombination accumulated across diverse accessions, but it may fail to detect rare alleles and is susceptible to population-structure confounding that generates false associations [4–6]. This limitation is particularly pronounced when panels are assembled from highly diverse, unadapted germplasm, where spurious associations can arise that reflect broad population differentiation rather than causal variation relevant to the defined geographic or agro-ecological target region. Systematically integrating both approaches through dual-platform verification could improve MQTL positional confidence and resolution simultaneously, facilitating efficient marker development, selection, and candidate gene identification [4, 6]. A further aspect absent from most existing maize meta-analyses is the systematic differentiation between broadly adaptive loci conferring consistent effects across environments and those providing environment-specific advantages (under drought, heat, or low-nitrogen stress). This information could have critical implications for deploying MQTLs strategically across diverse production systems and breeding programs targeting contrasting target environments [9, 39–40]. As climate variability intensifies globally, understanding which loci provide stable genetic benefits across environments versus environment-specific adaptive advantages becomes increasingly important for developing simultaneously productive and resilient crop varieties [41–44].

The availability of high-quality reference genome assemblies [45–47], comprehensive gene expression atlases covering diverse tissues and developmental stages [48–49], gene interaction networks enabling network-based candidate prioritization [50], and comparative genomics resources enabling cross-cereal translational analysis [47] now creates opportunities for meta-QTL analysis with substantially enhanced biological interpretation, extending well beyond positional refinement to systematic functional candidate gene prioritization, network-based evidence integration, and cross-species functional inference. We present a comprehensive multi-trait meta-analysis spanning 1995–2025 that integrates 2,701 QTLs from 304 independent studies across five trait categories: grain yield and components; plant development and architecture; plant physiology and stress adaptation; grain quality and nutritional composition; and disease and pest resistance. Our specific objectives were to: (i) consolidate QTLs into refined high-confidence meta-QTLs through dual-platform verification integrating linkage and association mapping signals; (ii) identify multi-trait genomic hotspots and classify them into functionally relevant categories that directly inform breeding strategy; (iii) classify the environmental context of MQTLs based on the environmental distribution of contributing QTLs to provide deployment guidance; (iv) systematically prioritize candidate genes within hotspot intervals using network-based analysis; and (v) assess cross-cereal syntenic conservation across rice, sorghum, wheat, and barley to enable translational breeding applications and cross-species marker transfer.

## Methods

### Data collection and literature mining

A systematic literature search across Google Scholar, Scopus, and PubMed was used to identify maize QTL mapping and GWAS studies published between 1995 and 2025. Search terms combined the keywords “maize”, “corn”, and “Zea mays” with “QTL”, “quantitative trait locus”, “GWAS”, and “genome-wide association”, together with trait-specific terms for each of the five targeted trait categories. The reference lists of major reviews, meta-analyses, and benchmark mapping studies were additionally screened to recover papers that automated database searches might have missed; the full list of contributing source studies is provided in Additional file 1 (sheet S2). A study was retained only when it satisfied four criteria: (i) it reported original QTL or GWAS data with sufficient positional information for projection onto a consensus map; (ii) it described clearly defined trait measurements obtained under consistent phenotyping protocols; (iii) it reported mapping population size and type, enabling confidence-interval (CI) estimation; and (iv) it validated the QTL or SNP–trait association statistically through permutation testing, LOD thresholding, or false-discovery-rate (FDR) control.

### QTL database construction and standardization

Each QTL was cataloged with its chromosome assignment, CI boundaries, peak-marker position, flanking-marker identities, LOD score or p-value, percentage of phenotypic variance explained (PVE), additive effect direction and magnitude, dominance effect (where reported), population type and size, trait name and measurement method, environmental conditions and stress treatment, and source publication. Trait nomenclature across the 304 source studies was harmonized by mapping each reported trait to the Crop Ontology (CO) vocabulary and assigning it to one of five functional categories: grain yield and components; plant development and architecture; plant physiology and stress adaptation; grain quality and nutritional composition; and disease and pest resistance. QTL positions reported under older marker systems were updated to current map coordinates using MaizeGDB marker correspondence tables and physical-position data. For the 6.9% of entries lacking a reported LOD score, a default value of 2.5 was assigned, consistent with established meta-QTL protocols [51–52]. Where CIs were not provided in the original study, they were estimated using population-type-specific formulas validated for each mapping design [53–54]: Throughout, PVE values are reported as member-MQTL PVE from the present meta-analysis unless explicitly labeled as single-study (individual biparental or association) estimates; single-study values represent upper-bound effect sizes subject to Beavis-effect inflation and are not directly comparable to member-MQTL PVE.

CI (backcross / F) = 530 / (N × r²)

CI (recombinant inbred lines) = 163 / (N × r²)

CI (doubled haploid) = 287 / (N × r²) where N is the mapping population size and r² is the proportion of phenotypic variance explained by the QTL. QTLs lacking both r² and CI information were excluded, because both parameters are required for precision-weighted projection.

### Consensus map construction and QTL projection

All QTLs were projected onto the IBM2 2008 Neighbors consensus genetic map, accessed through MaizeGDB. This map was selected because it has the broadest marker overlap with the populations, marker systems, and genetic backgrounds represented across three decades of published maize QTL studies, and because it has been used consistently throughout the published crop meta-QTL literature [15, 17, 20–21, 24, 52, 55]. SNP physical positions were converted to genetic coordinates using MaizeGDB marker correspondence tables. Where studies reported only flanking markers without explicit peak positions, those flanking markers were placed on the consensus framework to define CI boundaries, and peak positions were estimated by midpoint interpolation. Of the 2,701 compiled QTLs, 2,518 (93.2%) were successfully projected onto the consensus map; 183 (6.8%) were excluded because of insufficient or ambiguous positional information. The excluded QTLs were proportionally distributed across trait categories and chromosomes, confirming the absence of systematic projection bias that could skew trait-category representation in downstream analyses.

### Meta-QTL analysis using BioMercator V4.2.3

Meta-analysis was performed separately for each of the five trait categories using BioMercator V4.2.3 [15]. BioMercator estimates MQTL positions within a Gaussian mixture framework in which each contributing QTL is weighted inversely by the width of its confidence interval, so that more precisely mapped QTLs from larger populations exert greater influence on the consensus position. Specifically, each QTL i received a weight w□□ 1 / CI□². The optimal number of MQTLs per chromosome was selected using the Akaike Information Criterion (AIC) [56–57], which trades off goodness of fit against the number of parameters to identify the most parsimonious clustering solution at each chromosomal position. Physical positions of the identified MQTLs were projected onto the B73 RefGen_v5 assembly (Zm-B73-REFERENCE-NAM-5.0) by linear interpolation between the nearest flanking IBM2 2008 Neighbors markers, following the protocol of Zhao et al. [52] and Wang et al. [55]. Because the compiled studies originally reported positions on multiple B73 assemblies (RefGen_v1–v5), all coordinates were harmonized to RefGen_v5 prior to interpolation; positions that could not be assigned unambiguously to v5 were re-anchored by flanking-marker identity. All annotated protein-coding genes within each MQTL physical interval were retrieved via the qTeller comparative gene-expression module on MaizeGDB and cross-referenced against gene-model annotations in Ensembl Plants (https://plants.ensembl.org) to ensure comprehensive interval gene catalogs [52, 58].

### Dual-platform verification through GWAS co-localization

Significant marker–trait associations (MTAs meeting each study’s genome-wide significance or Bonferroni threshold) were collected from 89 published GWAS studies, yielding 855 SNP–trait associations spanning all five targeted trait categories. The physical positions of all GWAS-MTAs were converted to a uniform B73 RefGen_v5 coordinate system using the Assembly Converter tool in Ensembl Plants to ensure cross-study positional comparability. An MQTL was counted as dual-platform supported through GWAS co-localization when its physical interval fell within ±1 Mb of at least one significant GWAS-MTA [52].

### Hotspot identification and candidate gene prioritization

Genomic regions of interest were identified under three predefined criteria that capture complementary breeding-relevant features: (i) multi-trait hotspots, defined as three or more MQTLs from distinct trait categories within a 5 Mb window; (ii) multi-population loci, defined as a single MQTL supported by five or more independent mapping populations from different genetic backgrounds; and (iii) major-effect loci, defined as an MQTL with member PVE >20% (the top 5% by effect size across the full MQTL catalog). Criterion (i) is an enrichment criterion and was tested for statistical significance against a gene-density-weighted null model that excluded pericentromeric regions (±20 Mb from annotated centromeres), using permutation-based FDR correction (10,000 iterations); criteria (ii) and (iii) are single-locus threshold selections and are not enrichment tests. The criterion or criteria satisfied by each of the 23 hotspots are reported in Additional file 1 (sheet S4, Criteria_Met column), and the genome-organization enrichment statistic reported in the Results is restricted to the multi-trait [criterion (i)] hotspots to which the permutation test applies. The expected number of MQTLs per 20 Mb window was calculated from the total euchromatic genome length remaining after pericentromeric exclusion, scaled by the genome-wide MQTL density.

Candidate gene prioritization within hotspot intervals employed three complementary evidence layers. First, network-based ranking used MaizeNet hub scores, which integrate co-expression data, protein–protein interaction networks, and curated functional annotations [50], to identify genes with the strongest overall biological connectivity and the most coherent functional profiles relative to each hotspot’s associated traits. Second, homology-based mining used BLASTp searches against a curated set of functional genes from maize and other cereal species, applying a top-priority threshold of E-value < 1×10□□ and alignment score >100 to identify high-confidence functional homologs. Third, cross-cereal orthologs were called by reciprocal best-hit BLASTp against the rice, sorghum, wheat, and barley proteomes, retaining pairs above 70% amino-acid identity, with sensitivity analyses also conducted at 60% and 50% thresholds. Synteny conservation across the cereal genomes was assessed using SynMap2 within the CoGe comparative genomics platform [58].

### Statistical quality control

MQTL positional stability was assessed by bootstrap resampling (1,000 iterations, with a random 20% of contributing QTLs withheld without replacement in each iteration). Three stability thresholds were defined: stringent (>90% detection rate across iterations, positional SD <3 Mb), standard (>80%, SD <5 Mb), and permissive (>70%, SD <10 Mb); the standard threshold served as the primary criterion for designating an MQTL as positionally stable. The environmental context of each MQTL was classified descriptively by tabulating the proportion of its contributing QTLs detected under optimal growing conditions versus abiotic or biotic stress, as recorded in the original mapping studies. On this basis, each MQTL was annotated as broadly adaptive (≥2 contributing QTLs from optimal conditions and ≥1 from stress conditions), stress-specific (contributing QTLs predominantly from stress conditions), or unclassified (insufficient environmental coverage for confident assignment). BioMercator’s precision-weighted averaging provides partial structural mitigation of effect-size inflation by down-weighting QTLs from smaller populations with wide confidence intervals. All statistical analyses were performed in R v4.2.0 [59], with linear mixed-effects modeling via lme4 [60].

## Results

### Comprehensive QTL integration and chromosomal distribution

Literature mining spanning 1995–2025 yielded 2,701 QTLs from 304 independent studies encompassing diverse germplasm backgrounds (Table 1; Fig. 1; Additional file 2: Fig. S1). The represented germplasm included temperate × temperate crosses (42.3%), tropical × temperate crosses (31.8%), tropical × tropical crosses (18.4%), and diversity panels (7.5%) [61], providing broad coverage of the allelic diversity present in cultivated maize and ensuring that the identified meta-QTLs are not unduly influenced by any single heterotic group or geographic germplasm pool. The QTLs were distributed across the five trait categories as follows: GYC (1,142 QTLs; 42.3%), PDA (678 QTLs; 25.1%), PHS (514 QTLs; 19.0%), GQN (312 QTLs; 11.5%), and DPR (106 QTLs; 3.9%); category counts sum to 2,752 rather than 2,701 because QTLs reported for multiple traits are tallied in each relevant category, so the sum exceeds the number of unique QTLs. Notably, 68% of QTLs were reported after 2010, coinciding with the widespread adoption of high-density SNP arrays [62–63] and successive B73 genome releases [45–47] and expanded developmental transcriptome resources [64] that substantially improved mapping resolution and cross-study comparability.

**Fig. 1.**
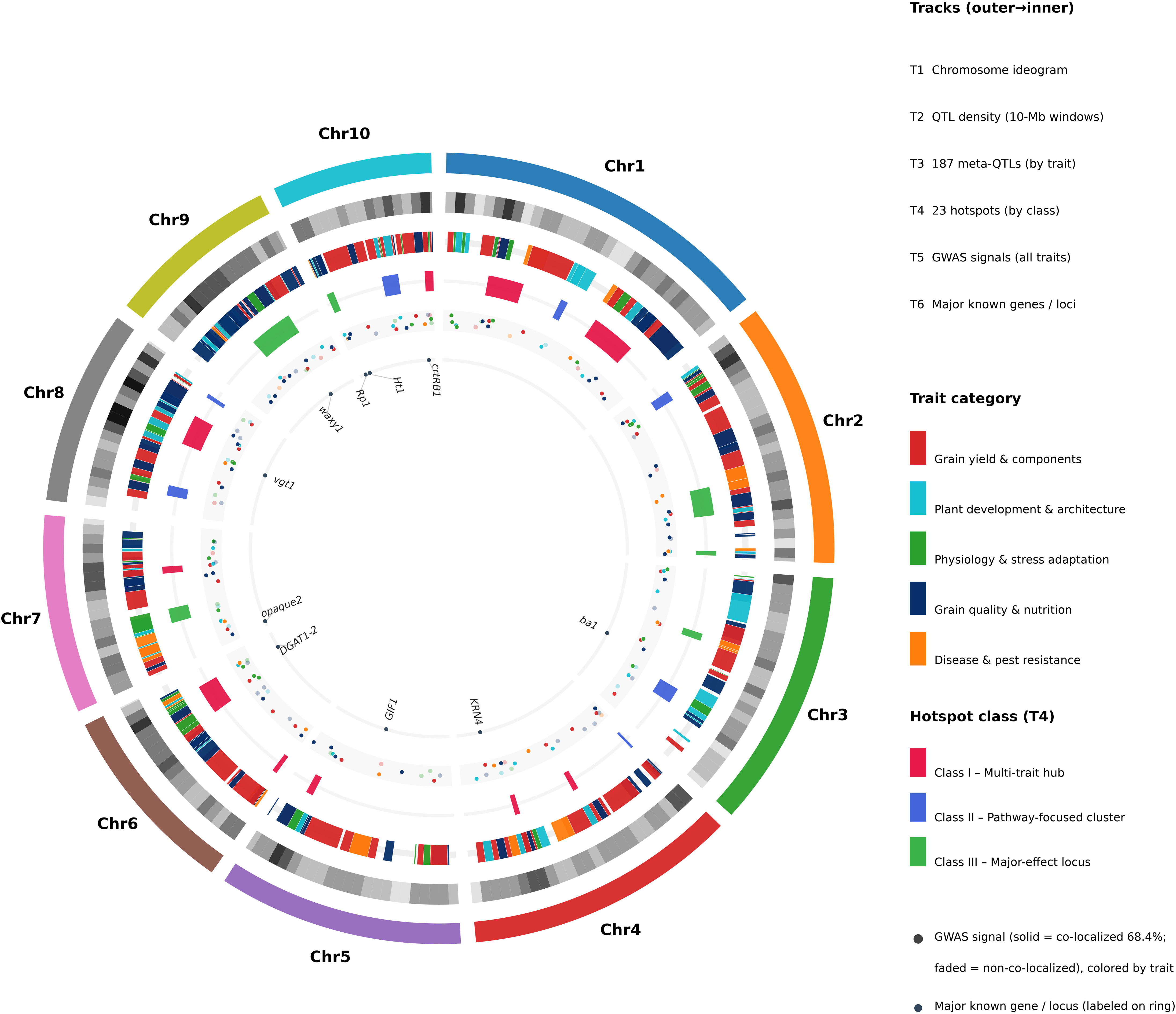
Circular genome map of meta-QTL distribution, genomic hotspots, and GWAS validation across the maize genome. Circular visualization integrating 187 meta-QTLs, 23 genomic hotspots, and genome-wide association signals across all ten maize chromosomes (B73 RefGen_v5). Six equally spaced concentric tracks display complementary layers (outermost to innermost). Track 1, chromosome ideogram with arc length proportional to physical chromosome size (≈150–307 Mb); chromosome identity is labeled at each arc midpoint. Track 2, gray-intensity meta-QTL density heatmap in 10 Mb sliding windows (darker = higher density). Track 3, the 187 high-confidence meta-QTLs shown as a single band at their consensus positions, colored by trait category: red (grain yield and components), cyan (plant development and architecture), green (physiology and stress adaptation), dark blue (grain quality and nutritional composition), and orange (disease and pest resistance); arc width reflects each meta-QTL consensus physical interval (clipped to chromosome bounds; genetic-map CI reduced 4.0-fold relative to source QTLs). Track 4, the 23 genomic hotspots (Table 2 core spans) colored by functional class: Class I multi-trait hubs (red), Class II pathway-focused clusters (blue), and Class III major-effect loci (green); these regions harbor 131 meta-QTLs (70.1%). Track 5, GWAS signals shown as a band spanning all trait categories, colored as in Track 3 and positioned at meta-QTL summits; co-localized signals (68.4%) are drawn at full opacity and non-co-localized signals (31.6%) are faded. Track 6 (innermost), major known genes/loci labeled directly on the ring with a marker at each genomic position: waxy1, opaque2, crtRB1, Rp1, Ht1, vgt1, KRN4, GIF1, ba1, and DGAT1-2. Abbreviations: CI, confidence interval; GWAS, genome-wide association study; Mb, megabase pairs; MQTL, meta-quantitative trait locus; QTL, quantitative trait locus.

**Table 1.**
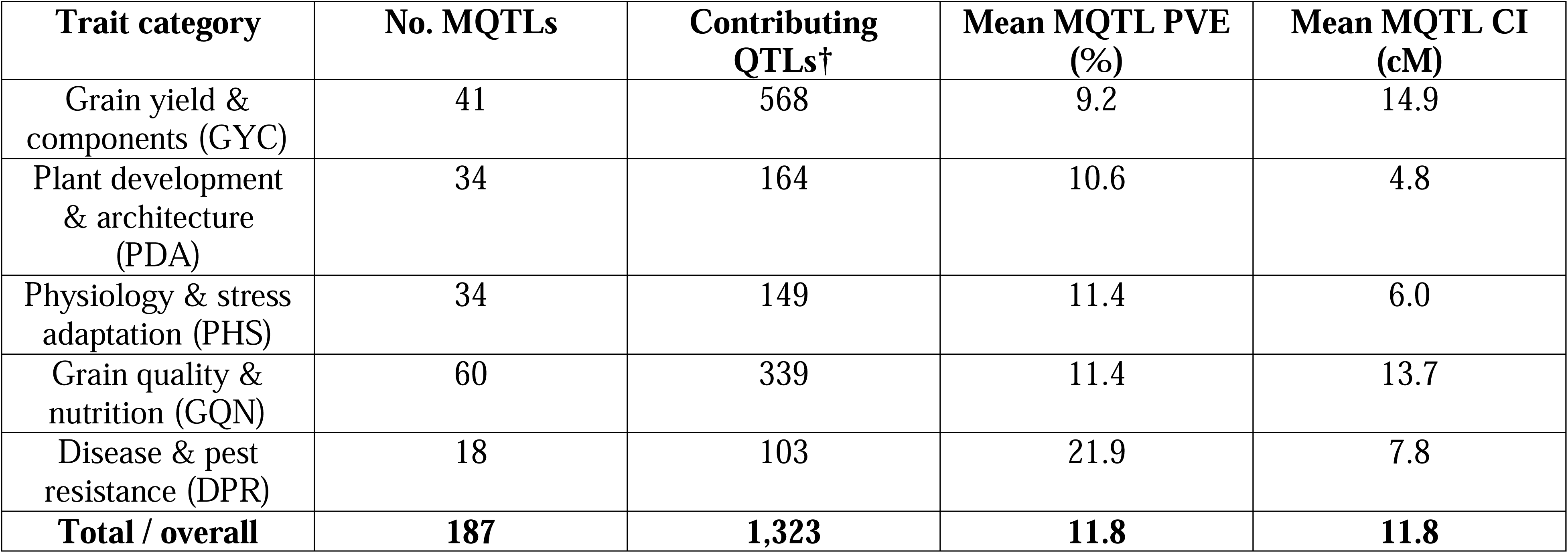
Summary of compiled QTLs and meta-QTL refinement by trait category. *Number of high-confidence meta-QTLs (MQTLs), contributing QTLs (the subset of the 2,518 projected QTLs that clustered into MQTLs; note this is fewer than the 2,518 total projected), mean MQTL phenotypic variance explained (PVE), and mean MQTL genetic-map confidence-interval (CI) width (cM) per trait category. Full per-MQTL data: Additional file 1 (sheets S1 and S2)*. *†“Contributing QTLs” counts the projected QTLs that were consolidated into the 187 MQTLs (column total 1,323); this is a subset of the 2,518 QTLs successfully projected onto the consensus map, because singleton projected QTLs that did not cluster into a meta-QTL are not counted here. (Per-category contributing-QTL counts are the sums of No_Initial_QTLs across each category’s MQTLs in Additional file 1.)*

**Table 2.**
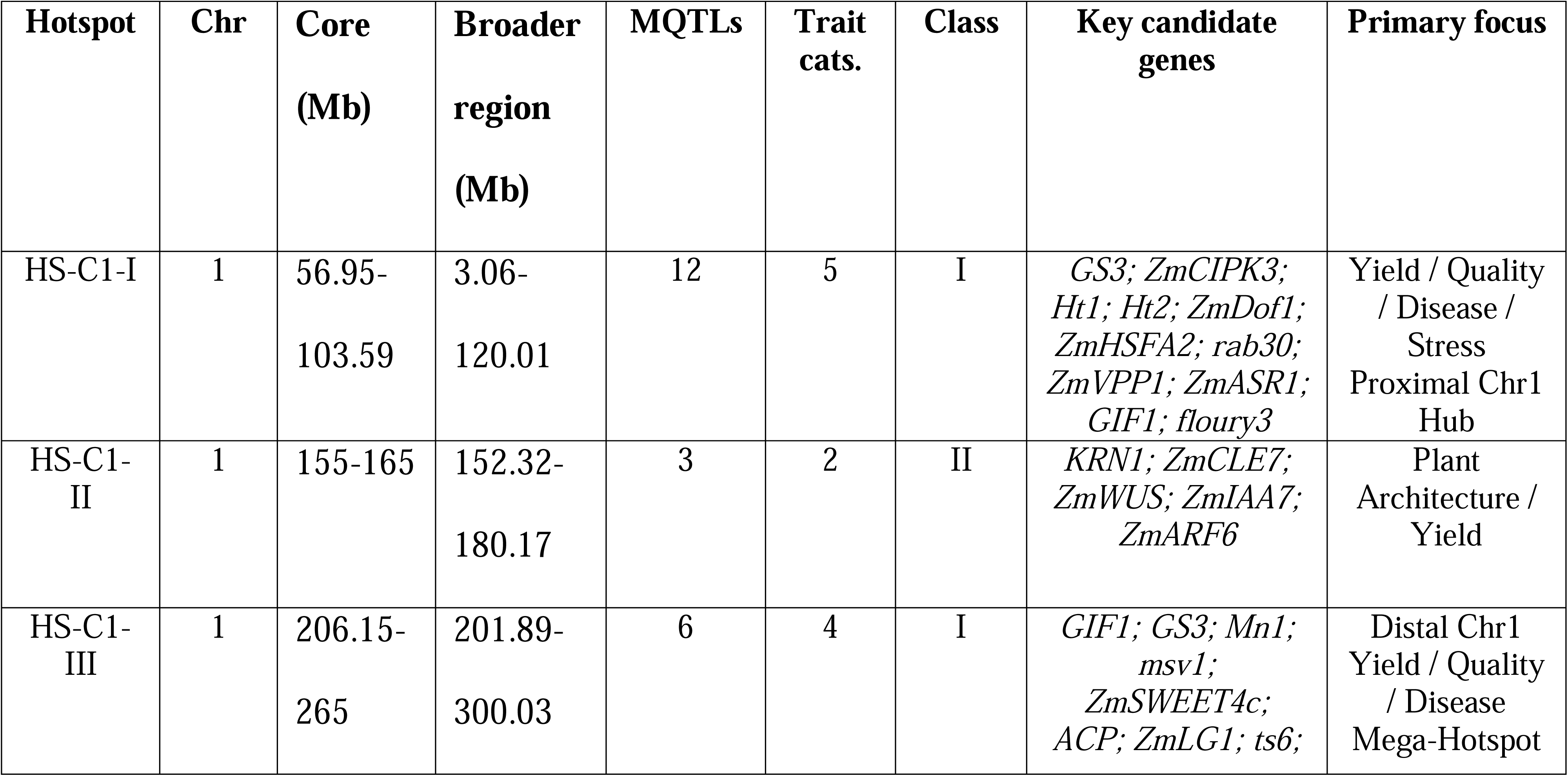
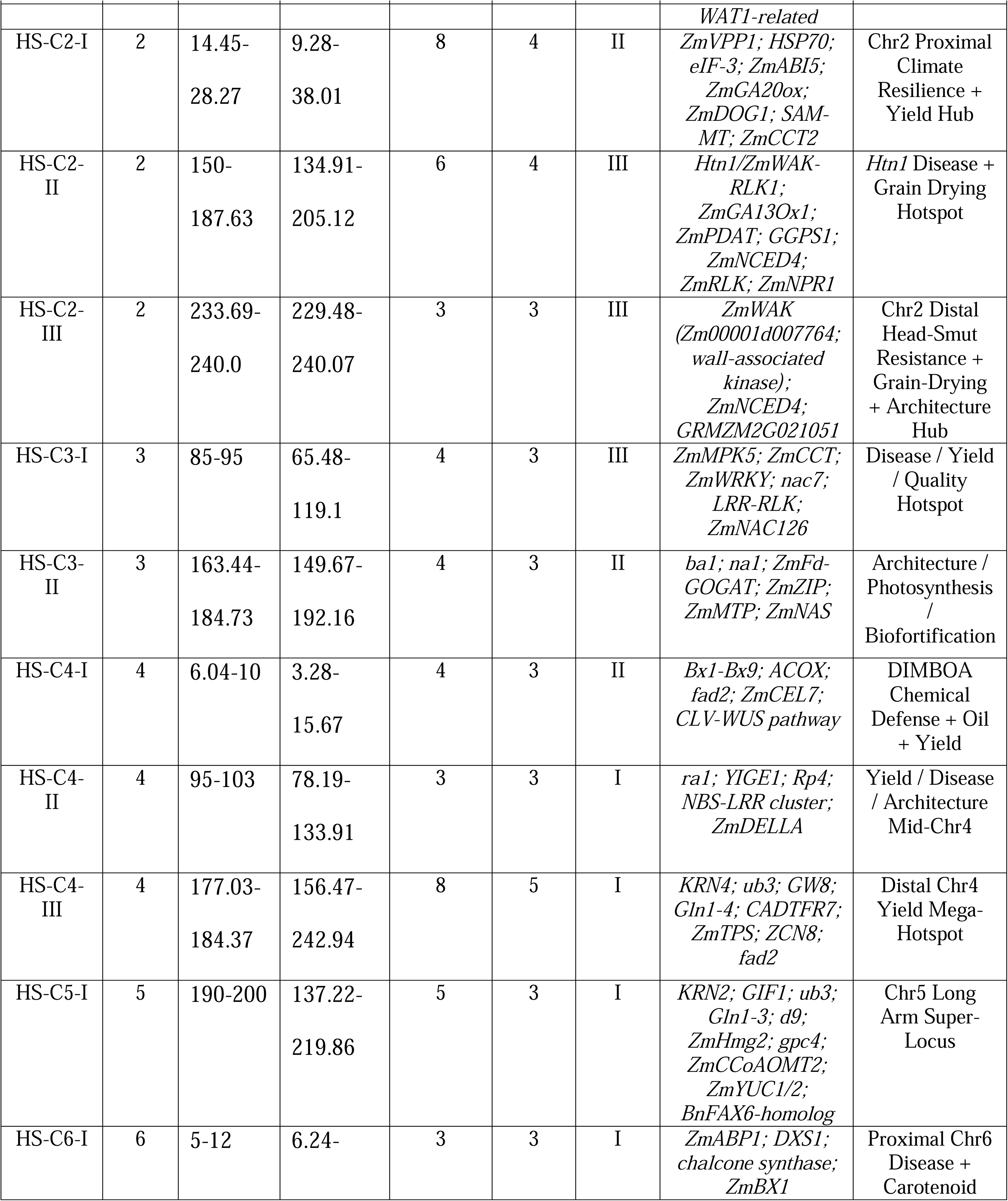
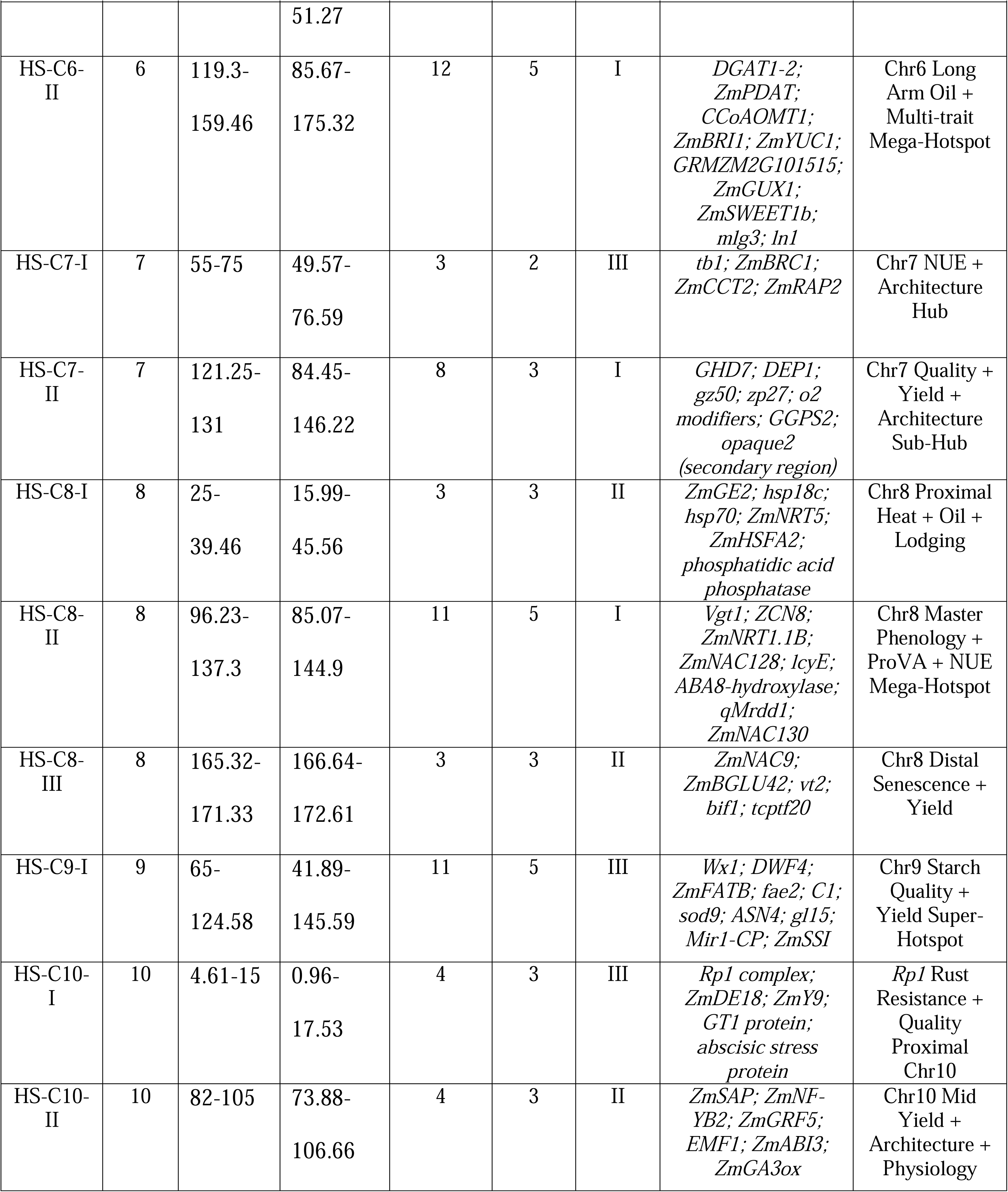
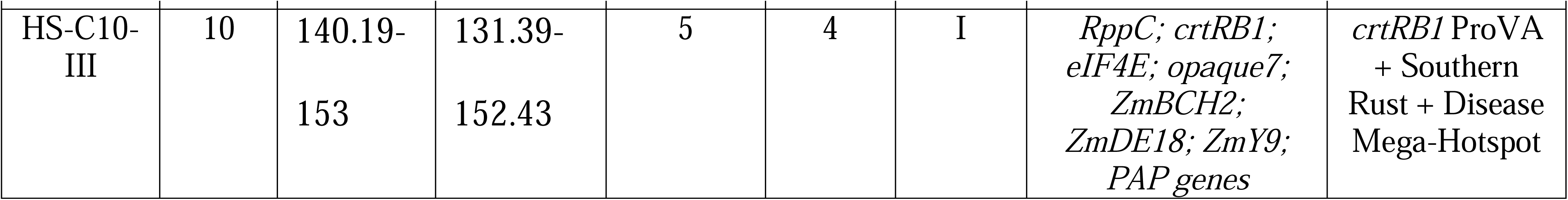
Genomic hotspots: coordinates, contributing MQTLs, trait categories, classification, and key candidate genes. *Hotspot coordinates (B73 RefGen_v5), number of contributing MQTLs and trait categories, and functional class assigned by an ordered, data-driven rule (I multi-trait hub, three or more substantive trait categories with no category contributing 50% or more of member-MQTL PVE; II pathway-focused cluster, one dominant pathway with maximum member-MQTL PVE below 28%; III major-effect locus, one dominant pathway with a member-MQTL PVE of 28% or higher), key candidate genes, and primary breeding focus. Class counts: I = 10, II = 7, III = 6. Full composition: Additional file 1 (sheet S4). Column of contributing-MQTL counts sums to 133 because two MQTLs (MQTL-GY1-4 and MQTL-DR4-2) each contribute to two overlapping hotspots; 131 distinct MQTLs (70.1% of 187) are located within hotspots*.

| Hotspot | Chr | Core<br>(Mb) | Broader<br>region<br>(Mb) | MQTLs | Trait<br>cats. | Class | Key candidate<br>genes | Primary focus |
| --- | --- | --- | --- | --- | --- | --- | --- | --- |
| HS-C1-I | 1 | 56.95-103.59 | 3.06-120.01 | 12 | 5 | I | GS3; ZmCIPK3;<br>Ht1; Ht2; ZmDof1;<br>ZmHSFA2; rab30;<br>ZmVPP1; ZmASR1;<br>GIF1; floury3 | Yield / Quality<br>/ Disease /<br>Stress<br>Proximal Chr1<br>Hub |
| HS-C1-II | 1 | 155-165 | 152.32-180.17 | 3 | 2 | II | KRN1; ZmCLE7;<br>ZmWUS; ZmIAA7;<br>ZmARF6 | Plant<br>Architecture /<br>Yield |
| HS-C1-III | 1 | 206.15-265 | 201.89-300.03 | 6 | 4 | I | GIF1; GS3; Mn1;<br>msv1;<br>ZmSWEET4c;<br>ACP; ZmLG1; ts6; | Distal Chr1<br>Yield / Quality<br>/ Disease<br>Mega-Hotspot |

| Hotspot | Chr | Core (Mb) | Broader region (Mb) | MQTLs | Trait cats. | Class | Key candidate genes | Primary focus |
| --- | --- | --- | --- | --- | --- | --- | --- | --- |
|  |  |  |  |  |  |  | *WAT1-related* |  |
| HS-C2-I | 2 | 14.45-28.27 | 9.28-38.01 | 8 | 4 | II | *ZmVPP1; HSP70; eIF-3; ZmABI5; ZmGA20ox; ZmDOG1; SAM-MT; ZmCCT2* | Chr2 Proximal Climate Resilience + Yield Hub |
| HS-C2-II | 2 | 150-187.63 | 134.91-205.12 | 6 | 4 | III | *Htn1/ZmWAK-RLK1; ZmGA13Ox1; ZmPDAT; GGPS1; ZmNCED4; ZmRLK; ZmNPR1* | *Htn1* Disease + Grain Drying Hotspot |
| HS-C2-III | 2 | 233.69-240.0 | 229.48-240.07 | 3 | 3 | III | *ZmWAK (Zm00001d007764; wall-associated kinase); ZmNCED4; GRMZM2G021051* | Chr2 Distal Head-Smut Resistance + Grain-Drying + Architecture Hub |
| HS-C3-I | 3 | 85-95 | 65.48-119.1 | 4 | 3 | III | *ZmMPK5; ZmCCT; ZmWRKY; nac7; LRR-RLK; ZmNAC126* | Disease / Yield / Quality Hotspot |
| HS-C3-II | 3 | 163.44-184.73 | 149.67-192.16 | 4 | 3 | II | *ba1; na1; ZmFd-GOGAT; ZmZIP; ZmMTP; ZmNAS* | Architecture / Photosynthesis / Biofortification |
| HS-C4-I | 4 | 6.04-10 | 3.28-15.67 | 4 | 3 | II | *Bx1-Bx9; ACOX; fad2; ZmCEL7; CLV-WUS pathway* | DIMBOA Chemical Defense + Oil + Yield |
| HS-C4-II | 4 | 95-103 | 78.19-133.91 | 3 | 3 | I | *ra1; YIGE1; Rp4; NBS-LRR cluster; ZmDELLA* | Yield / Disease / Architecture Mid-Chr4 |
| HS-C4-III | 4 | 177.03-184.37 | 156.47-242.94 | 8 | 5 | I | *KRN4; ub3; GW8; Gln1-4; CADTFR7; ZmTPS; ZCN8; fad2* | Distal Chr4 Yield Mega-Hotspot |
| HS-C5-I | 5 | 190-200 | 137.22-219.86 | 5 | 3 | I | *KRN2; GIF1; ub3; Gln1-3; d9; ZmHmg2; gpc4; ZmCCoAOMT2; ZmYUC1/2; BnFAX6-homolog* | Chr5 Long Arm Super-Locus |
| HS-C6-I | 6 | 5-12 | 6.24- | 3 | 3 | I | *ZmABP1; DXS1; chalcone synthase; ZmBX1* | Proximal Chr6 Disease + Carotenoid |
|  |  |  | 51.27 |  |  |  |  |  |
| HS-C6-II | 6 | 119.3-159.46 | 85.67-175.32 | 12 | 5 | I | *DGAT1-2*;<br>*ZmPDAT*;<br>*CCoAOMT1*;<br>*ZmBRI1*; *ZmYUC1*;<br>*GRMZM2G101515*;<br>*ZmGUX1*;<br>*ZmSWEET1b*;<br>*mlg3*; *ln1* | Chr6 Long<br>Arm Oil +<br>Multi-trait<br>Mega-Hotspot |
| HS-C7-I | 7 | 55-75 | 49.57-76.59 | 3 | 2 | III | *tb1*; *ZmBRC1*;<br>*ZmCCT2*; *ZmRAP2* | Chr7 NUE +<br>Architecture<br>Hub |
| HS-C7-II | 7 | 121.25-131 | 84.45-146.22 | 8 | 3 | I | *GHD7*; *DEP1*;<br>*gz50*; *zp27*; *o2*<br>*modifiers*; *GGPS2*;<br>*opaque2*<br>(secondary region) | Chr7 Quality +<br>Yield +<br>Architecture<br>Sub-Hub |
| HS-C8-I | 8 | 25-39.46 | 15.99-45.56 | 3 | 3 | II | *ZmGE2*; *hsp18c*;<br>*hsp70*; *ZmNRT5*;<br>*ZmHSFA2*;<br>*phosphatidic acid*<br>*phosphatase* | Chr8 Proximal<br>Heat + Oil +<br>Lodging |
| HS-C8-II | 8 | 96.23-137.3 | 85.07-144.9 | 11 | 5 | I | *Vgt1*; *ZCN8*;<br>*ZmNRT1.1B*;<br>*ZmNAC128*; *lcyE*;<br>*ABA8-hydroxylase*;<br>*qMrdd1*;<br>*ZmNAC130* | Chr8 Master<br>Phenology +<br>ProVA + NUE<br>Mega-Hotspot |
| HS-C8-III | 8 | 165.32-171.33 | 166.64-172.61 | 3 | 3 | II | *ZmNAC9*;<br>*ZmBGLU42*; *vt2*;<br>*bif1*; *tcptf20* | Chr8 Distal<br>Senescence +<br>Yield |
| HS-C9-I | 9 | 65-124.58 | 41.89-145.59 | 11 | 5 | III | *Wx1*; *DWF4*;<br>*ZmFATB*; *fae2*; *C1*;<br>*sod9*; *ASN4*; *gl15*;<br>*Mir1-CP*; *ZmSSI* | Chr9 Starch<br>Quality +<br>Yield Super-Hotspot |
| HS-C10-I | 10 | 4.61-15 | 0.96-17.53 | 4 | 3 | III | *Rp1 complex*;<br>*ZmDE18*; *ZmY9*;<br>*GT1 protein*;<br>*abscisic stress*<br>*protein* | *Rp1* Rust<br>Resistance +<br>Quality<br>Proximal<br>Chr10 |
| HS-C10-II | 10 | 82-105 | 73.88-106.66 | 4 | 3 | II | *ZmSAP*; *ZmNF-YB2*; *ZmGRF5*;<br>*EMF1*; *ZmABI3*;<br>*ZmGA3ox* | Chr10 Mid<br>Yield +<br>Architecture +<br>Physiology |
| HS-C10-III | 10 | 140.19-153 | 131.39-152.43 | 5 | 4 | I | *RppC*; *crtRB1*;<br>*eIF4E*; *opaque7*;<br>*ZmBCH2*;<br>*ZmDE18*; *ZmY9*;<br>*PAP genes* | *crtRB1* ProVA<br>+ Southern<br>Rust + Disease<br>Mega-Hotspot |

QTL distribution varied significantly across chromosomes. Chromosome 1 harbored the most QTLs (348; 12.9%), followed by chromosome 4 (312; 11.5%). When normalized by physical length, chromosomes 9 and 10 exhibited the highest densities, at 2.38 and 2.39 QTLs per 10 Mb, respectively, suggesting particularly rich genetic variation controlling agronomic traits in these regions (Additional file 3: Fig. S2). Approximately 38.2% of QTLs localized to pericentromeric regions (±20 Mb from annotated centromeres), reflecting suppressed recombination and reduced mapping resolution in these chromosomal domains [54]. Environmental classification revealed that 1,661 QTLs (61.5%) were identified under optimal growing conditions, 704 QTLs (26.1%) under abiotic stress, and 336 QTLs (12.4%) under biotic stress. Among the 704 abiotic-stress QTLs, drought-responsive loci accounted for the largest fraction (47.3%), followed by low-nitrogen stress (22.1%), heat stress (18.9%), and combined or other abiotic stresses (11.7%), reflecting the historical emphasis of the maize genetics community on drought as the primary abiotic constraint to productivity.

### QTL effect sizes vary among trait categories

PVE differed substantially among trait categories (Additional file 1: sheet S1). At the category level, mean member-MQTL PVE was highest for disease and pest resistance (21.9%; Table 1), reflecting the major-gene architecture typical of R-gene-mediated resistance together with the small number of resistance MQTLs (18), which raises the category mean. At the individual-locus level, however, the largest single effects were concentrated in grain quality and nutritional composition, where individual biochemical pathway genes can exert substantial quantitative effects on composition; 3.8% of grain-quality QTLs exceeded 20% PVE, including starch-biosynthesis genes (wx1, ae1, su1; member-MQTL PVE range 7.4–52.0%) [65–66], the endosperm protein regulator opaque2 (25.0% member-MQTL PVE) [67], and the carotenoid-biosynthesis gene crtRB1 (14.0% member-MQTL PVE) [68]. Plant development and architecture QTLs included several major-effect loci: vgt1 (up to ∼54% PVE for flowering time in individual biparental populations [69], an upper-bound single-study estimate subject to Beavis-effect inflation; meta-QTL member PVE 10.3%, MQTL-PS8-2, consistent with the highly polygenic architecture of maize flowering time), the brachytic2/dwarf3 plant-height locus [70], and barren stalk1 (24.3% PVE) [71]. In contrast, grain yield and components QTLs exhibited smaller individual effects, with none of the grain-yield MQTLs exceeding 20% PVE, consistent with the highly polygenic architecture of yield traits, which reflects many small-effect loci distributed across the genome [3, 72].

Physiological and stress-adaptation QTLs showed conditional effect amplification under stress relative to optimal conditions. For drought-responsive QTLs evaluated under contrasting water regimes within the same mapping populations, average PVE increased from 3.8% under well-watered conditions to 13.4% under severe drought, a 3.5-fold amplification (paired t-test, P < 0.001) [20, 30]. This conditional amplification has important implications for the choice of genomic-selection training environments and for the strategic deployment of stress-specific versus broadly adaptive loci in breeding programs.

### Integration and refinement of meta-QTLs

Of the 2,701 compiled QTLs, 2,518 (93.2%) were successfully projected onto the IBM2 2008 Neighbors consensus genetic map, while 183 (6.8%) were excluded because of insufficient or ambiguous positional information; these exclusions were proportionally distributed across trait categories and chromosomes. BioMercator V4.2.3 consolidated the 2,518 projected QTLs into 187 high-confidence MQTLs, each consolidating on average 7.1 contributing QTLs (range 2–28; contributing QTLs = the 1,323 projected QTLs that clustered into MQTLs, Table 1; MQTLs supported by only two contributing QTLs were retained [52, 55]). Because meta-analysis operates on the genetic map, confidence-interval refinement is quantified on a genetic-distance basis: the mean 95% CI contracted 4.0-fold, from 47.6 cM across cM-reported contributing QTLs to 11.8 cM in the MQTLs (median 29.1 to 8.2 cM; 75% reduction; Table 1; Fig. 2), narrowing the candidate-gene search space across all trait categories. This reduction is comparable to the 2.5– to 5.9-fold reductions reported for single-trait maize meta-analyses of yield, abiotic-stress, disease-resistance, and morphological traits [20–24], and is achieved here across five heterogeneous trait categories rather than a single trait. Physical (Mb) coordinates are reported to anchor each MQTL to the reference genome and to enable candidate-gene retrieval; because each MQTL is the consensus envelope of on average 7.1 clustered source QTLs, its physical span (mean 16.4 Mb after quality control; Fig. 2) is necessarily broader than that of an individual source QTL and is therefore not used as a measure of positional refinement, which the maize genetic-to-physical map (with pericentromeric ratios exceeding 10 Mb cM ¹) would otherwise distort.

**Fig. 2.**
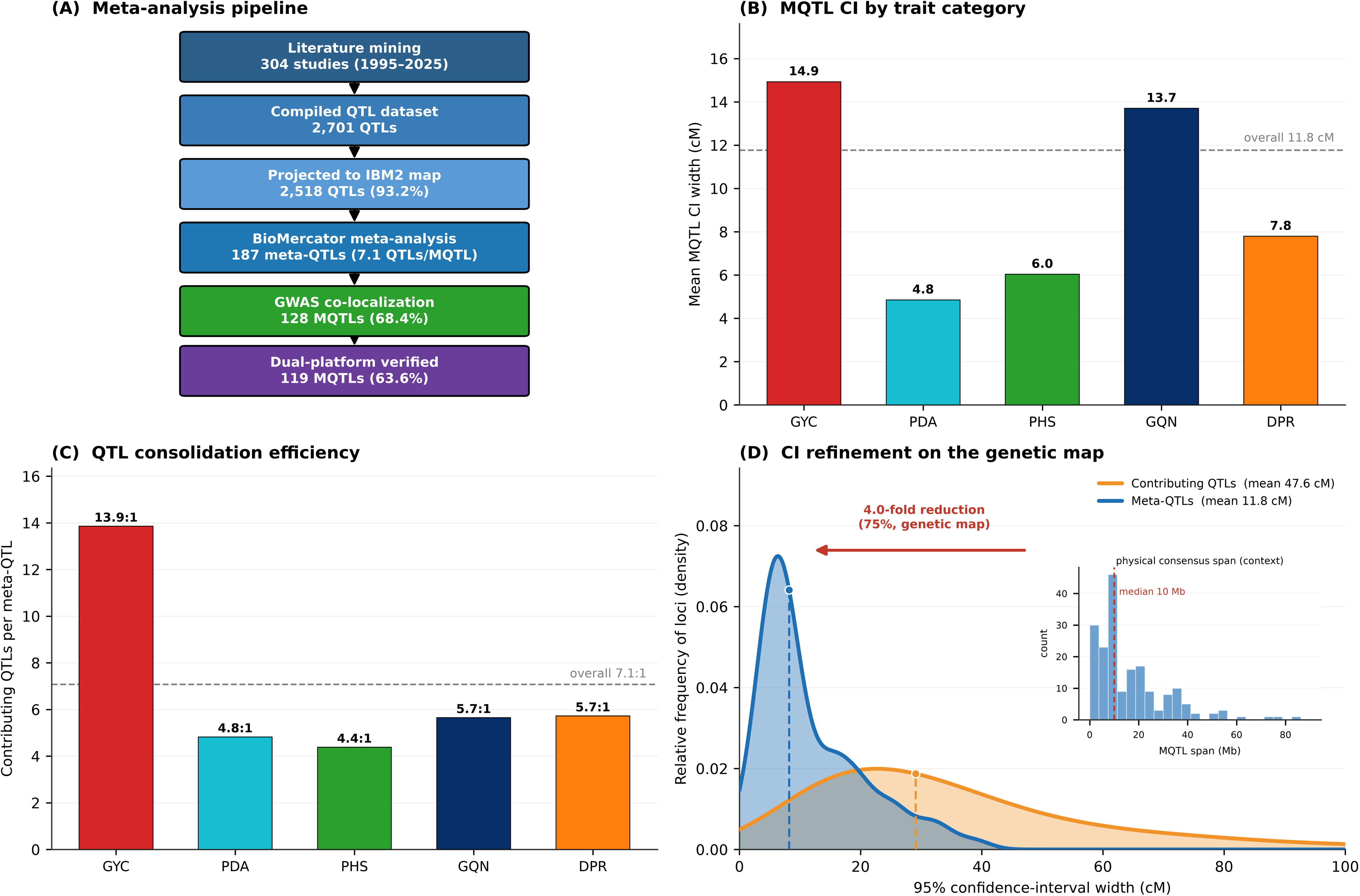
Meta-QTL integration and dual-platform verification. (A) Pipeline: 2,701 QTLs from 304 studies were projected onto the IBM2 consensus map (2,518; 93.2%) and integrated into 187 meta-QTLs (overall 13.5 projected QTLs per meta-QTL; equivalently 7.1 contributing QTLs per meta-QTL); 128 (68.4%) showed GWAS co-localization and 119 (63.6%) were dual-platform verified. (B) GWAS co-localization rate by trait category; circles are scaled to the number of meta-QTLs per category (grain quality and nutritional composition 85.2%, n = 60; plant development and architecture 83.3%, n = 34; grain yield and components 77.5%, n = 41; physiology and stress adaptation 69.0%, n = 34; disease and pest resistance 57.9%, n = 18); dashed line, overall 68.4%. (C) Consolidation efficiency (contributing QTLs per meta-QTL): grain yield and components 13.9:1, grain quality and nutritional composition 5.7:1, disease and pest resistance 5.7:1, plant development and architecture 4.8:1, physiology and stress adaptation 4.4:1; dashed line, overall 7.1:1. (D) Confidence-interval refinement on the genetic map (mean 47.6 → 11.8 cM, 4.0-fold/75% reduction; median 29.1 → 8.2 cM); the physical (Mb) distribution of MQTL consensus spans is shown for genomic context. MQTL, meta-QTL; GWAS, genome-wide association study.

Interval refinement varied considerably among trait categories, reflecting differences in the number and precision of contributing studies. Grain-yield and components MQTLs showed the greatest improvement (62% reduction; 32.5 to 12.4 Mb), driven by the large number of contributing yield studies. Disease-resistance MQTLs also refined substantially (31% reduction; 21.6 to 15.0 Mb), while physiology and stress-adaptation MQTLs were already narrow before meta-analysis and changed little (6.7 to 6.4 Mb). Across the full set, 4.3% of MQTLs refined below 1 Mb, intervals sufficiently narrow to directly inform gene-level functional studies. The most extensively supported MQTLs across all categories included MQTL-GY4-5, harboring *KRN4* (25 contributing QTLs); MQTL-GQ9-4, harboring the waxy/amylose starch-quality gene *wx1* (*waxy1*) (8 contributing QTLs); the flowering-time locus MQTL-PS8-2, harboring *vgt1* (6 contributing QTLs); and MQTL-GY5-2, harboring *GIF1* (24 contributing QTLs).

Bootstrap resampling (1,000 iterations, 20% QTL exclusion per iteration) confirmed that 170 of 187 MQTLs (90.9%) were positionally consistent when any 20% subset of contributing QTLs was randomly withheld, indicating that the identified MQTL positions are not unduly driven by any single subset of the compiled literature. Stability was consistent across all five trait categories: GQN (90.7%), GYC (90.0%), PDA (88.9%), DPR (84.2%), and PHS (83.3%). MQTLs supported by ≥10 underlying QTLs showed near-universal stability (63 of 64; 98.4%), confirming that multi-study support is the most reliable predictor of positional robustness.

### Dual-platform verification through GWAS co-localization

Integration of 855 significant marker–trait associations from 89 GWAS studies provided GWAS co-localization support for 128 of 187 MQTLs (68.4%), defined as co-localization within ±1 Mb of at least one GWAS signal meeting genome-wide significance thresholds (Additional file 1: sheet S3). Of these 128 GWAS-supported MQTLs, 119 additionally satisfied the stricter dual-platform criterion requiring concordant evidence from both linkage mapping (LOD ≥ 3.0) and association mapping, representing 63.6% of all 187 MQTLs. This cross-platform concordance substantially increases confidence that these MQTL positions reflect genuine genetic signals rather than population-specific mapping artifacts. Functional confirmation at individual loci nevertheless requires targeted fine-mapping to causal genes and rigorous experimental validation through genetic transformation, CRISPR editing, or near-isogenic line comparisons.

GWAS co-localization rates varied meaningfully by trait category, reflecting differences in the available GWAS literature and in the genetic architectures of different trait classes: GQN achieved the highest rate (85.2%), followed by PDA (83.3%), GYC (77.5%), PHS (69.0%), and DPR (57.9%). For GWAS-supported MQTLs, association mapping narrowed genomic intervals to an average width of 420 kb containing approximately 18 annotated genes, a substantial improvement over the linkage-only median CI of 12.6 Mb containing hundreds of candidate genes. Notable examples of GWAS-enabled refinement included MQTL-GY9-3 (reduced from 12.8 Mb to 20 kb) [73], *waxy1* (22.6 Mb to 0.68 Mb), *Dwarf3* (18.7 Mb to 0.18 Mb), and *opaque2* (18.9 Mb to 0.84 Mb), demonstrating the power of dual-platform integration for achieving gene-level resolution in favorable cases. Biological-consistency assessment confirmed that, after coordinate correction, 118 of 119 independently characterized maize genes with known functions (99.2%; Table 3) mapped within an MQTL confidence interval, the sole exception (*Htn1*) being assigned by chromosome-bin to MQTL-DR8-1; characterized genes showed preferential concentration in refined intervals below the median CI of 12.6 Mb relative to the 50% expected under a random distribution (P < 0.001). This figure reflects the internal consistency of the gene-to-MQTL assignments rather than independent validation, since the characterized-gene set and the MQTL intervals were derived from the same underlying catalog.

**Table 3.**
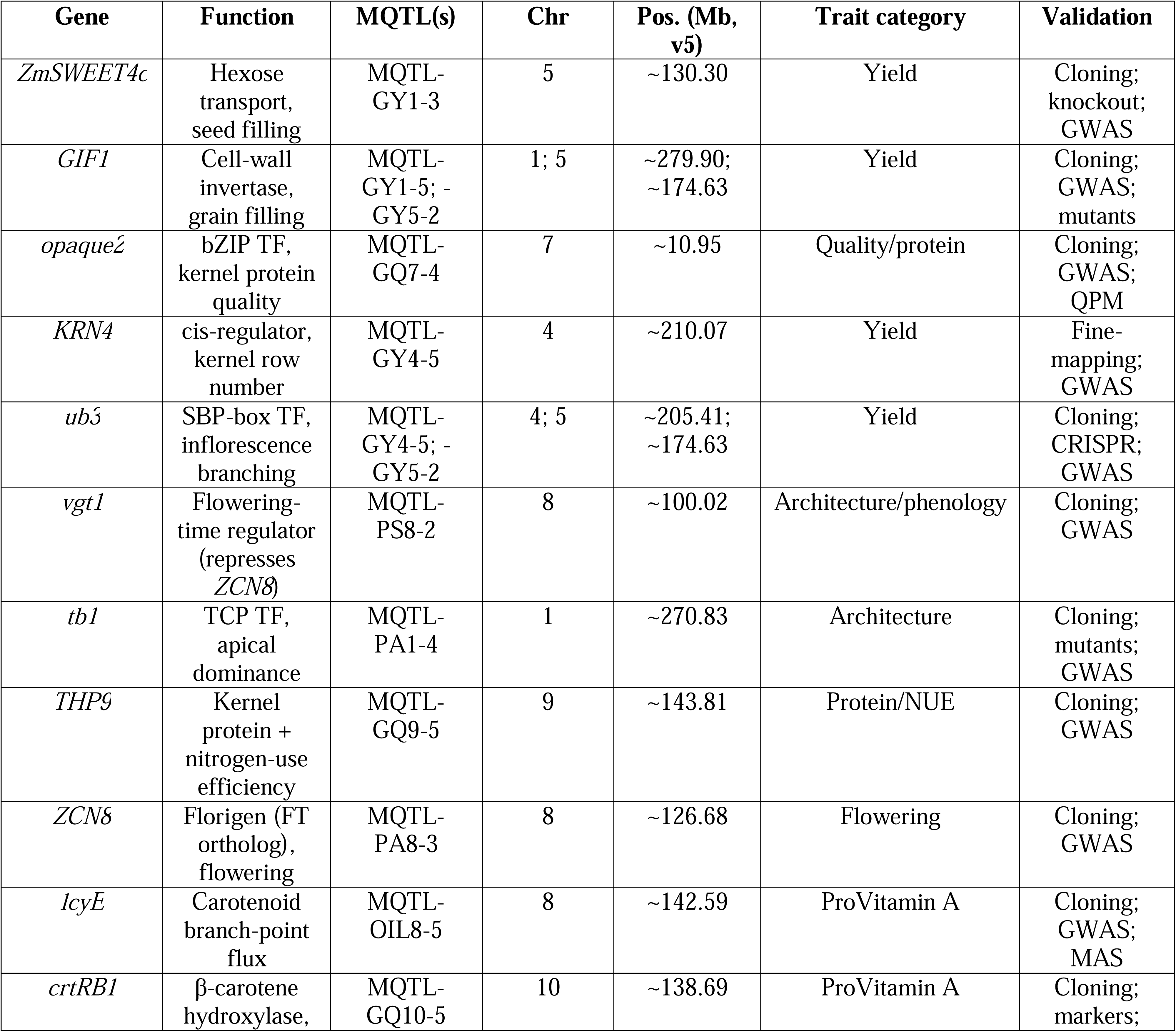
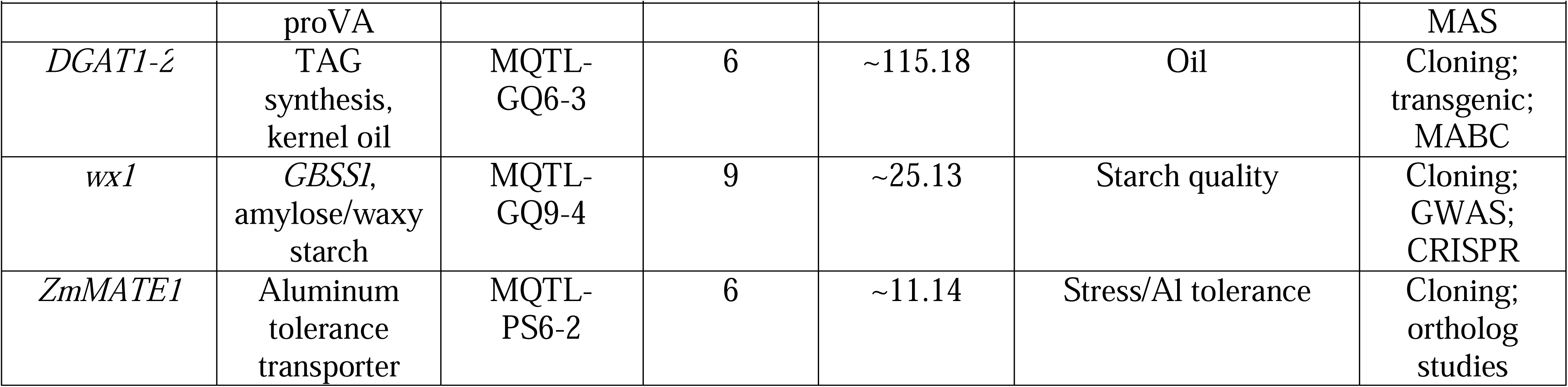
Representative priority genes within genomic hotspots (validation status indicated per gene). *Gene name, function, associated MQTL(s), physical position (B73 RefGen_v5), trait category and validation for an illustrative subset; the full validated set (119 genes) with references and confidence levels is provided as Additional file 1 (sheet S5)*.

The meta-QTLs successfully captured major domestication genes (*tb1*, *tga1*, *ba1*), quality determinants (*wx1*, *ae1*, *su1*, *opaque2*, *crtRB1*), and grain-yield regulators (*KRN1*, *KRN4*, *GIF1*), validating the biological relevance of the identified genomic regions across diverse functional gene classes.

### Genomic hotspots and multi-trait co-localization

Systematic analysis identified 23 genomic hotspots meeting the three predefined criteria (Table 2). These hotspots occupy a core footprint of approximately 478 Mb (the summed consensus-overlap intervals of member MQTLs; “Core (Mb)” column, Table 2), corresponding to 25.6% of the ∼1,866 Mb euchromatic genome, yet harbor 131 of 187 MQTLs (70.1%), a concentration significantly exceeding chance expectations from permutation testing (P < 0.001, FDR-corrected). Measured instead as the broader consensus regions encompassing all flanking member-QTL confidence intervals (“Broader region (Mb)” column, Table 2), the 23 hotspots span approximately 1,179 Mb; the two columns bound the hotspot footprint at its dense consensus core and its maximal flanking extent, respectively. The remaining 56 non-hotspot MQTLs were distributed across the remaining ∼1,388 Mb of euchromatic sequence. Hotspot MQTLs showed significantly higher mean PVE than non-hotspot MQTLs (15.6% vs. 6.5%; Welch’s t-test, P < 0.001), confirming that genomic hotspots harbor disproportionately larger genetic effects and are not simply an artifact of regional MQTL clustering within gene-dense regions. The multi-trait architecture was robust to category subsampling, with all 10 Class I hubs recovered in >90% of subsampling iterations and environment-dependent co-localization patterns preserved in >95%, confirming that the observed hotspot structure is not driven by any single trait category’s contribution to the compiled database.

Multi-trait co-localization analysis, defined as MQTLs from different trait categories located within 5 Mb of each other and validated by sensitivity analyses at 2.5 Mb and 10 Mb windows revealed environment-dependent patterns of genomic co-localization. Among MQTLs predominantly supported by optimal-condition QTLs, grain-yield MQTLs co-localized frequently with plant development and architecture MQTLs (62.5%) and grain-quality MQTLs (34.2%), consistent with source–sink relationships and shared developmental pathways. Among MQTLs predominantly supported by drought-condition QTLs, co-localization shifted markedly toward physiological and stress-adaptation loci (71.4%), particularly those affecting anthesis– silking interval and leaf senescence, processes that become rate-limiting specifically under osmotic stress. Under low-nitrogen conditions, yield MQTLs showed increased co-localization with nitrogen-use-efficiency and remobilization loci (58.3%), highlighting the metabolic overlap between nitrogen assimilation and grain filling. These shifting patterns suggest that different gene sets control yield performance across contrasting environments, although they also partly reflect the environmental distribution of published mapping studies [9, 39]. Disease and pest resistance MQTLs showed limited co-localization with other categories (<15% with any single category), suggesting that resistance loci can generally be selected with minimal expected effects on agronomic performance under disease-free conditions. A small number of cross-category MQTL overlaps within single refined intervals of <500 kb were observed, several of them within Class I multi-trait hubs (Table 2); resolving whether these reflect genuine biological pleiotropy or tight physical linkage of functionally distinct genes will require targeted fine-mapping and CRISPR-based functional dissection.

### Classification of genomic hotspots

The 23 genomic hotspots were classified into three categories based on the patterns of multi-trait co-localization and effect sizes observed across contributing MQTLs (Table 2).

Class I: Multi-trait hubs (10 hotspots; 43.5%). These regions carry member meta-QTLs spanning three or more substantive trait categories with no single category contributing 50% or more of the summed member-MQTL PVE, and they represent the most immediate opportunities for simultaneous multi-trait improvement through coordinated selection (Fig. 3). HS-C6-II on chromosome 6 (125.88-166.32 Mb) is the most functionally diverse hotspot in the dataset, harboring 12 MQTLs spanning five trait categories and including the globally deployed oil-biosynthesis gene DGAT1-2 [74], the aluminum-tolerance gene ZmMATE2 [75], the brassinosteroid-signaling receptor ZmBRI1 [76], Fusarium resistance loci, and very-long-chain fatty-acid composition QTLs, a functionally diverse assembly of overlapping loci that collectively make this interval one of the highest-priority regions for multi-trait breeding investment. Two hotspots that harbor a canonical large-effect gene nonetheless qualify as Class I hubs because, at the meta-QTL level, their member effects are distributed across three substantive trait categories with no single category dominating variance: the vgt1/ZCN8 flowering-time region (HS-C8-II), which co-localizes developmental, grain-quality, and disease-resistance meta-QTLs alongside the provitamin-A gene lcyE [81], and the crtRB1 provitamin-A region (HS-C10-III), which co-localizes carotenoid-quality, yield, physiology, and disease-resistance meta-QTLs including southern-rust resistance (RppC) and eIF4E-mediated recessive potyvirus resistance [68]. Among co-localizing MQTL pairs within Class I hubs for which the favorable-allele direction was reported in the contributing studies, the majority showed concordant favorable-allele directions, indicating predominantly synergistic multi-trait relationships, while a minority showed discordant directions, indicating antagonistic relationships that require strategic management in breeding; a systematic, quantitative concordance analysis across all hubs would require complete, harmonized allelic-direction records and is identified here as a priority for follow-up work, such as accepting modest trade-offs in lower-priority traits to achieve gains in primary breeding targets.

**Fig. 3.**
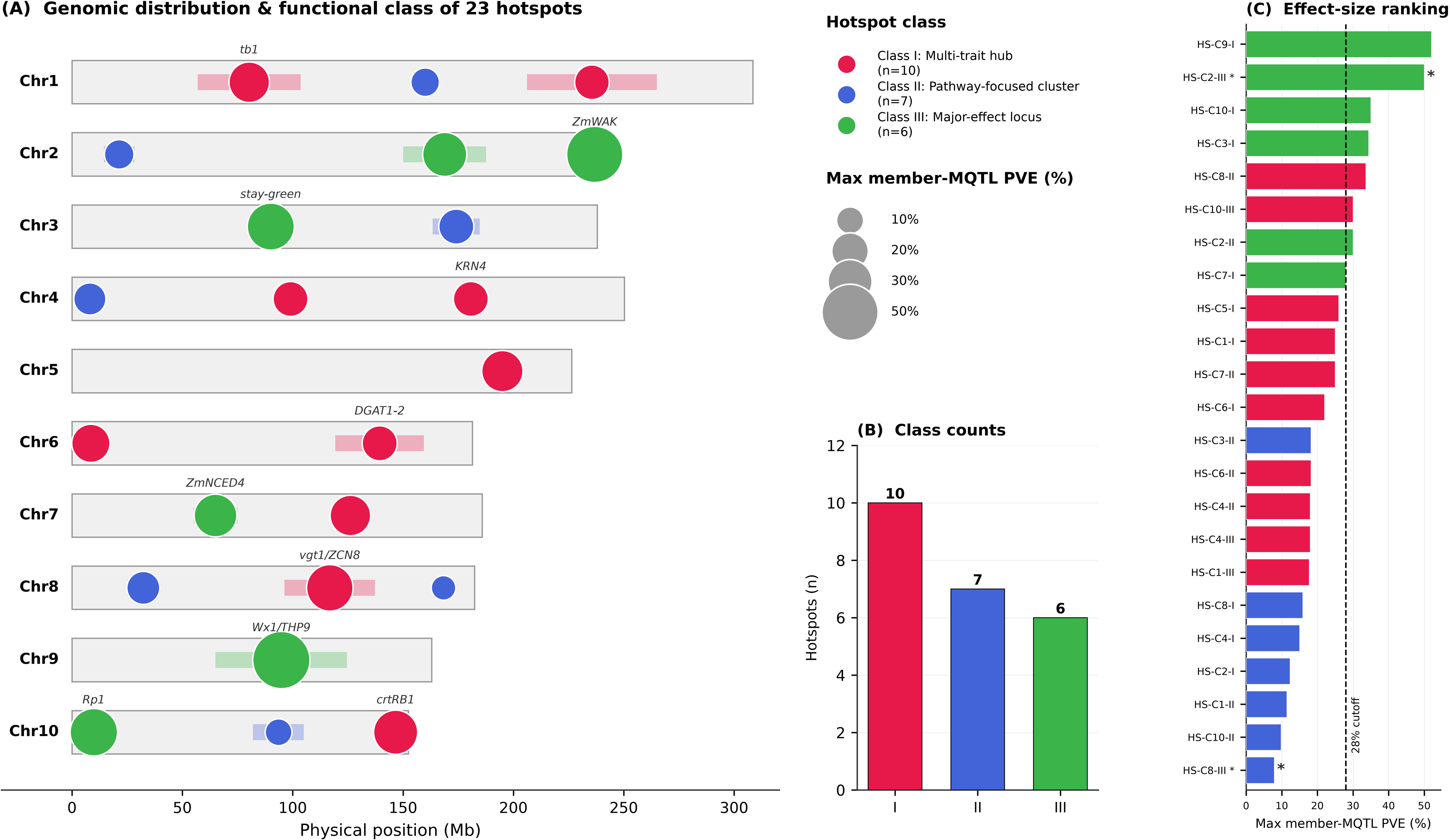
Genomic distribution and functional classification of the 23 multi-trait hotspots. (A) Physical positions of the 23 hotspots across the ten maize chromosomes (B73 RefGen_v5); each hotspot is drawn at its consensus-core interval, colored by functional class (Class I multi-trait hubs (red; n = 10), Class II pathway-focused clusters (blue; n = 7), and Class III major-effect loci (green; n = 6)), with marker size scaled to the maximum member-MQTL phenotypic variance explained (PVE) and key anchoring loci labeled (tb1, KRN4, DGAT1-2, vgt1/ZCN8, crtRB1, ZmWAK, stay-green, ZmNCED4, Wx1/THP9, Rp1). (B) Hotspot counts per functional class. (C) Effect-size ranking of the hotspots by maximum member-MQTL PVE, with the 28% Class III major-effect threshold marked; asterisks flag hotspots whose defining member meta-QTL rests on minimum study support. PVE, phenotypic variance explained; MQTL, meta-quantitative trait locus.

Class II: Pathway-focused clusters (7 hotspots; 30.4%). These regions are dominated by a single trait pathway, meaning that one trait category contributes the majority of member-MQTL PVE or that the hotspot spans no more than two substantive categories, while member effects remain moderate (maximum member-MQTL PVE below 28%). Minor meta-QTLs from other categories may co-localize, but the genetic architecture of the region is functionally centred on one pathway, offering opportunities for focused trait improvement with limited expected effects on other agronomic characteristics. The clearest example is the chromosome 8 kernel-yield cluster (HS-C8-I), in which 83.0% of the summed member-MQTL PVE is concentrated in a single trait pathway (Additional file 1: sheet S4), so focused improvement can proceed with minimal off-target co-localization; the cluster is anchored by the triple-validated ZmNRT5 lodging-resistance and yield-component loci [80]. Further examples include the chromosome 4 grain-quality and oil cluster (HS-C4-I), which carries the DIMBOA benzoxazinoid insect-defense and kernel-oil meta-QTLs, and plant-architecture-focused clusters on chromosomes 1, 3, and 10 (HS-C1-II, HS-C3-II, and HS-C10-II). In each, the dominant-pathway meta-QTLs can be selected with predictably limited trade-offs on non-target traits. Chromosome 9 additionally harbors ZmFatB (GRMZM5G829544), where a single G-nucleotide insertion truncates the acyl-ACP thioesterase and reduces palmitic acid and total saturated fatty acids, improving oil quality without lowering total oil content [79].

Class III: Major-effect loci (6 hotspots; 26.1%). These hotspots are pathway-focused but carry at least one member meta-QTL of large effect (member-MQTL PVE of 28% or higher), placing them among the strongest loci in their trait categories and distinguishing them from the moderate-effect clusters of Class II. Applying this quantitative criterion rather than a curated list of named genes identifies six loci. The chromosome 9 Wx1/THP9 starch-and-protein locus (HS-C9-I; 67.12-128.66 Mb; member-MQTL PVE up to 52.0%) is dominated by the starch-biosynthesis and quality-pathway genes Wx1, ae1, and su1 [65–66] together with THP9 (asparagine synthetase; chr9, B73 RefGen_v5), which markedly increases kernel protein content and nitrogen-use efficiency [82]. The head-smut resistance locus HS-C2-III (chr2 approximately 229.48-240.07 Mb; 50.0% member-MQTL PVE) is defined by ZmWAK and co-localizes internode-length and grain-drying physiology meta-QTLs. The Rp1 common-rust resistance locus (HS-C10-I; 35.0% member-MQTL PVE; 8 independent studies) is defined by the major-effect Rp1 R-gene complex. The chromosome 3 stay-green locus HS-C3-I (85.37-96.25 Mb; 34.4% member-MQTL PVE) carries senescence-delaying alleles at nac7 and ZmNAC126 that may extend photosynthetic duration, enhance nutrient remobilization efficiency, and improve grain nutritional quality through prolonged nitrogen accumulation [77–78]. A chromosome 2 disease-resistance locus (HS-C2-II; 30.0% member-MQTL PVE) and the chromosome 7 anthesis-silking-interval drought locus (HS-C7-I; 28.0% member-MQTL PVE; ZmNCED) complete the class. These loci are high-priority targets for marker-assisted selection and precision breeding, given their large and reproducibly detected effects across multiple independent studies and diverse germplasm backgrounds; the complete set of 13 MQTLs exceeding 20% member-MQTL PVE, led by wx1 (52.0%), is provided in Additional file 1: sheet S1.

### Environmental classification and candidate gene prioritization

Annotation of QTL environmental contexts identified three groups reflecting the distribution of published literature across environmental conditions. MQTLs predominantly supported by optimal-condition QTLs (broadly adaptive; 80 MQTLs; 42.8%), defined as those with ≥2 contributing QTLs from optimal-condition studies and ≥1 from stress-condition studies, were enriched for grain quality (81.0% of quality MQTLs) and plant development (68.2%) (Additional file 4: Fig. S3). Of these broadly adaptive MQTLs, 45 (56.3%) had contributing QTLs spanning three or more distinct optimal and two or more distinct stress contexts, suggesting broad adaptation across a wide range of growing conditions. MQTLs predominantly supported by stress-condition QTLs (stress-specific; 54 MQTLs; 28.9%), in which contributing QTLs were detected exclusively or predominantly under abiotic or biotic stress, included 25 drought-responsive, 12 low-nitrogen-responsive, 10 heat-responsive, and 7 biotic-stress-responsive loci, collectively covering all major stress categories relevant to maize production. The remaining 53 MQTLs (28.3%) had insufficient environmental coverage for confident assignment and were left unclassified. Average contributing-QTL PVE was significantly higher for stress-specific than for broadly adaptive MQTLs (12.8% vs. 7.4%). The drought-responsive ASI locus harboring ZmNCED4 exemplified environmental conditionality most strikingly, with contributing-QTL PVE increasing from 3.2% under well-watered conditions to 16.8% under severe drought, a 5.3-fold increase consistent with ABA-mediated pollen viability becoming rate-limiting under osmotic stress [20, 30] (Additional file 1: sheet S3).

Network-based prioritization using MaizeNet hub scores [50] and BLASTp homology against curated functional genes from maize and other cereal species markedly narrowed the candidate gene pools within all 23 hotspot intervals, with top-priority candidates showing strong concordance with known functional roles and significant enrichment of transcription factors, starch-biosynthesis enzymes, flowering-time regulators, senescence-related genes, and drought– and nitrogen-stress-response pathway members. Cross-cereal ortholog analysis (≥70% amino-acid identity; Table 4) identified prioritized candidates with confirmed orthologs in one or more of the comparison cereals, with conservation strongest for maize–rice and maize–sorghum gene pairs. Conservation was highest for developmental and metabolic genes, including Wx1/GBSSI, the florigen ZCN8, and ZmMATE2 [75], and lowest for pathogen-resistance genes, consistent with faster evolutionary turnover at host–pathogen coevolutionary interfaces. Seven hotspot regions contained top-priority candidates for which no confirmed functional counterparts were detected in the comparison cereals under the applied search parameters, including THP9 on chromosome 9 (within the MQTL-GQ9-5 protein-content interval), which encodes asparagine synthetase 4 (ZmASN4) and raises kernel protein content while improving nitrogen-use efficiency [82]; the highest-LOD ASI drought MQTL at HS-C10-II (LOD = 13.1), which lacked ortholog coverage for its key candidates [20]; and the Bx1–Bx9 benzoxazinoid cluster at HS-C4-I, which evolved in the Zea–Tripsacum clade after divergence from sorghum [83]. These intervals provide tractable entry points for NIL-based fine-mapping, CRISPR-mediated functional characterization, and synteny-guided gene discovery in related cereal species.

**Fig. 4.**
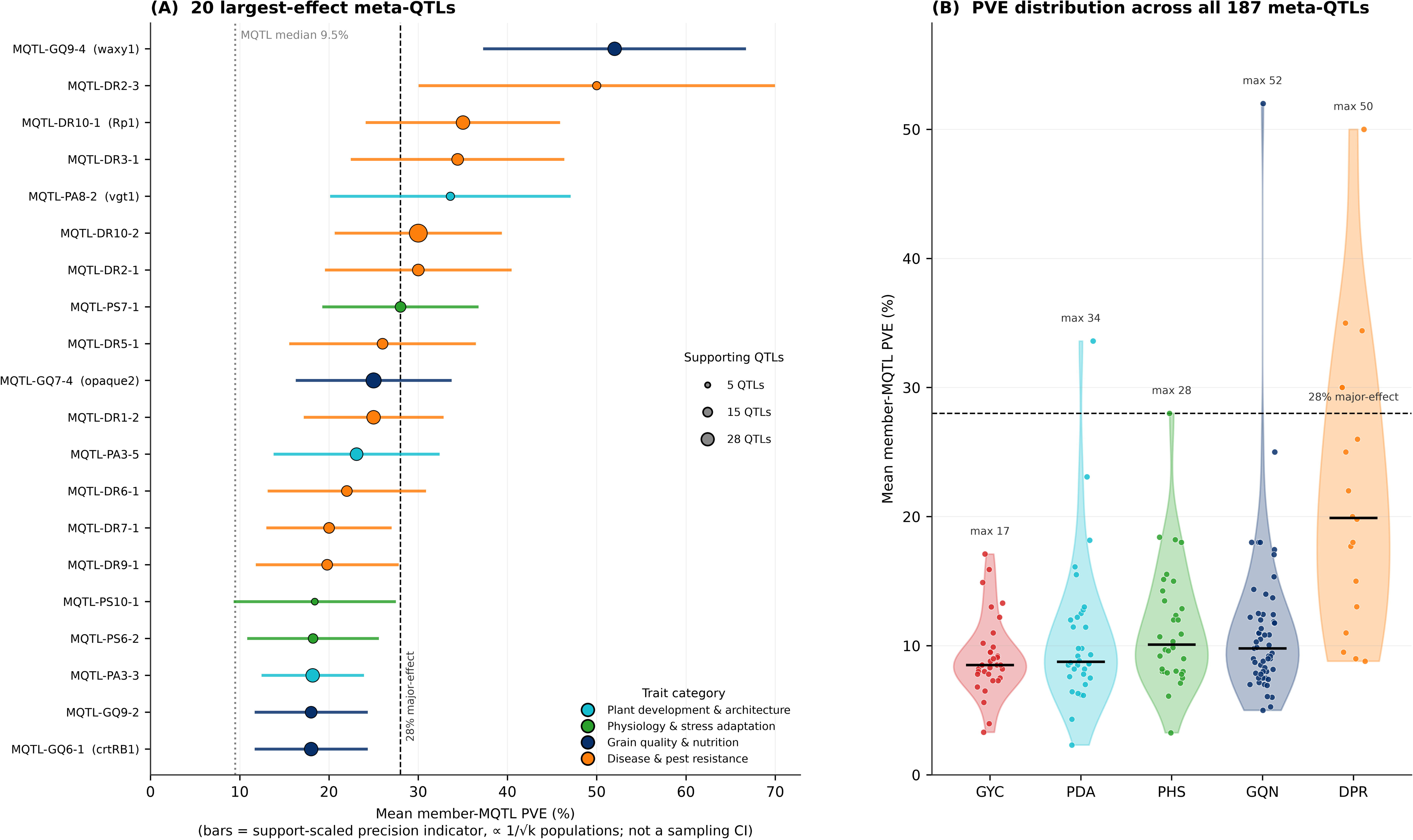
Effect-size architecture of the meta-QTL catalog. (A) Forest plot of the 20 largest-effect meta-QTLs ranked by mean member-MQTL phenotypic variance explained (PVE); points are colored by trait category and scaled to the number of supporting QTLs, and horizontal bars show a support-scaled precision indicator ( 1/√k populations; not a sampling confidence interval). Well-characterized grain-quality and developmental loci (waxy1, opaque2, crtRB1, vgt1) show the largest individual-locus effects; the 28% major-effect threshold and the genome-wide MQTL median (9.5%) are marked. (B) Distribution of mean member-MQTL PVE across all 187 meta-QTLs by trait category (GYC, grain yield and components; PDA, plant development and architecture; PHS, plant physiology and stress adaptation; GQN, grain quality and nutritional composition; DPR, disease and pest resistance); the per-category maximum is annotated and the 28% major-effect threshold is marked. Absolute PVE values reflect published estimates and are subject to Beavis-effect inflation. PVE, phenotypic variance explained; MQTL, meta-quantitative trait locus.

**Table 4.** Cross-cereal conserved genes and orthologs (validated and candidate). Maize candidate genes and their orthologs in rice, sorghum, wheat, and barley with BLASTp % amino-acid identity (best-hit model gene) and mean identity. Full ortholog table (102 records): Additional file 1 (sheet S6).

| Maize gene | Function | Rice (%ID) | Sorghum (%ID) | Wheat (%ID) | Barley (%ID) | Mean %ID |
| --- | --- | --- | --- | --- | --- | --- |
| *wx1* | GBSSI; amylose synthesis | 85 (*Os06g0133000*) | ortholog (*Sobic.010G022600*) | 85 (*Wx-A1/B1/D1*) | 85 (*HvGBSSI*) | 85 |
| *fad2* | $\Delta 12$-desaturase; oleic acid | 82 (*OsFAD2*) | 80 (*SbFAD2*) | 82 (*TaFAD2*) | ortholog (*HvFAD2*) | 81 |
| *ZCN8* | Florigen (FT); | 78 (*Hd3a*) | ortholog (*SbFT*) | 78 (*VRN3/TaFT*) | 79 (*HvFT1*) | 78 |
|  | flowering |  |  |  |  |  |
| *ZmGA20ox* | GA biosynthesis; plant height | 75 (*OsGA20ox2/SD1*) | ortholog (*SbGA20ox*) | 75 (*TaGA20ox*) | 78 (*HvGA20ox*) | 76 |
| *ZmGW2* | RING E3 ligase; grain size | 71 (*OsGW2*) | 69 (*SbGW2*) | 68 (*TaGW2*) | ortholog (*HvGW2*) | 69 |
| *Dwarf3* | GA/CYP pathway; plant height | 68 (*OsD3/CYP*) | ortholog (*SbCYP*) | 65 (*TaCYP*) | 68 (*HvCYP*) | 67 |
| *opaque2* | bZIP TF; kernel protein quality | 45 (*OsbZIP*) | 50 (*SbO2*) | 45 | - | 47 |

## Discussion

### Enhanced resolution through integrative meta-analysis

The dual-platform approach, leveraging the complementary strengths of biparental linkage mapping and association mapping in diverse germplasm panels, produced high-confidence, well-resolved meta-QTLs that substantially exceed the resolution achievable by either approach independently. GWAS co-localization support was obtained for 68.4% of MQTLs at the primary ±1 Mb threshold. This window is a support indicator rather than a positional-resolution metric, and a formal permutation-based enrichment test against a random-MTA null, together with a sensitivity analysis at narrower windows, would further quantify the strength of this support; nonetheless, co-localization for the majority of meta-QTLs is consistent with these positions representing genuine genetic signals rather than positional artifacts of individual mapping populations. Grain quality and nutritional composition achieved the highest dual-platform co-localization rate (85.2%), likely because large-effect biochemical pathway genes such as wx1, opaque2, and crtRB1 produce strong, consistent signals detectable across both mapping frameworks and diverse germplasm [25–26]. Confidence-interval refinement is reported on the genetic map (mean 47.6 → 11.8 cM; 75% reduction; Fig. 2D), consistent with the treatment of genetic distance as the valid refinement metric throughout this study. Physical intervals are provided only to anchor each meta-QTL to the reference genome for candidate retrieval and are not interpreted as a measure of positional refinement; where a co-localizing GWAS-MTA is available it provides an independent single-marker anchor within the meta-QTL interval that can guide subsequent fine-mapping, but it does not itself narrow the meta-QTL confidence interval. The 31.6% of MQTLs lacking GWAS co-localization support represent lower-confidence targets that warrant additional mapping effort before deployment; the reduced disease-resistance co-localization rate (57.9%) likely reflects the relative scarcity of published diversity-panel GWAS studies for resistance traits and the inherent difficulty of detecting race-specific resistance alleles in diverse panels where resistance-breaking pathogen races may be present [22–23]. The Beavis effect systematically inflates PVE estimates compiled across diverse studies and mapping populations [7–8]; meta-analytic precision-weighted averaging provides partial mitigation through down-weighting of QTLs from smaller populations, but absolute PVE values should be interpreted with caution, and comparative rankings among MQTLs are more reliable indicators of relative effect-size magnitude than absolute values.

Hotspot biology and candidate gene mechanisms

The 23 hotspots harbor 131 of 187 MQTLs (70.1%) within 478 Mb (25.6%) of the ∼1,866 Mb euchromatic genome (Fig. 3; Additional file 2: Fig. S1). Restricting the enrichment test to the multi-trait [criterion (i)] hotspots to which the permutation null applies, the concentration of multi-trait meta-QTLs within these regions, which was robust to trait-category subsampling, is consistent with non-random co-localization of multi-trait loci rather than a consequence of overall gene-density variation. The biological basis of synergistic multi-trait co-localization is illustrated most clearly by the chromosome 6 hub (HS-C6-II), where oil-biosynthesis (DGAT1-2), aluminum-tolerance (ZmMATE2), and disease-resistance meta-QTLs co-localize with no single pathway dominating variance, three agronomically beneficial effect classes achievable through coordinated selection at a single genomic region and representing a strong opportunity for efficient multi-trait improvement. The predominance of concordant favorable-allele directions among direction-resolvable co-localizing MQTL pairs indicates mostly synergistic relationships that breeders could exploit through coupling-phase selection, whereas the minority showing discordant directions identify regions where favorable alleles for one trait create unfavorable effects on others, requiring target-environment-specific prioritization, exploitation of tissue-specific expression variation to functionally uncouple antagonistic effects, or precision editing of regulatory sequences to achieve desired combinations [84–85]. The HS-C6-II multi-trait hub on chromosome 6 exemplifies the scale of potential simultaneous gains achievable at a single region: DGAT1-2 [74] driving oil-content improvement, ZmMATE2 [75] conferring aluminum tolerance on acid soils covering approximately 40-50% of tropical arable land, and Fusarium resistance loci offering pathogen protection, all within a 40 Mb interval that could in principle be targeted through a coordinated marker-assisted introgression.

A small number of cross-category MQTL overlaps within single refined intervals of <500 kb were observed, several of which fall within Class I multi-trait hubs; these provide stronger circumstantial evidence for biological pleiotropy or extremely tight linkage than broader hotspot-level co-localization patterns alone, although co-localization at this resolution still cannot distinguish a single pleiotropic gene from two tightly linked genes. Definitive resolution of the pleiotropy-versus-linkage question requires fine-mapping below 100 kb combined with CRISPR-based functional dissection of individual candidate genes, a distinction with direct and substantial practical consequences, since the pleiotropic effects of a shared gene cannot be separated by recombination, whereas tightly linked genes can potentially be decoupled through strategic crossing programs producing large segregating populations [84–85]. Pathway-focused clusters represent a contrasting and equally valuable architectural principle enabling focused trait improvement with substantially reduced trade-off risk: the chromosome 8 kernel-yield cluster (HS-C8-I) concentrates 83.0% of its summed member-MQTL PVE in a single trait pathway (Additional file 1: sheet S4, Dominant_PVE_share column), so focused improvement can proceed with minimal off-target co-localization and quality or yield gains can be captured efficiently without expected penalties on non-target traits [80]. Disease-resistance clusters are attractive candidates for concurrent resistance-gene pyramiding, and the near-neutral fitness effects of most resistance alleles in the absence of pathogen challenge [86–87] predict limited agronomic penalty under disease-free conditions; however, the coupling-versus repulsion-phase configuration of these linked resistance loci was not determined here and would need to be established from phased haplotype data in the target germplasm before pyramiding strategy is finalized. The functional classification of hotspots into these three strategically distinct categories thus provides breeders with an actionable decision framework: multi-trait hubs when simultaneous improvement of multiple traits is the primary objective and the program can invest in managing potential trade-offs; pathway-focused clusters when focused quality or resistance improvement with minimal unintended consequences is required; and major-effect loci when maximizing genetic gain for a single high-priority trait justifies large-scale marker-assisted selection.

### Environmental classification and deployment implications

MQTLs predominantly supported by optimal-condition QTLs (broadly adaptive; 42%) are enriched for grain quality (81%) and plant development (68%), suggesting that conserved metabolic enzymes and developmental regulators maintain consistent genetic effects across diverse environments. The higher PVE of stress-condition-supported MQTLs (12.8% vs. 7.4%) is consistent with stress conditions exposing rate-limiting genetic steps that are otherwise buffered by redundant pathways under favorable conditions [20, 88]. The *ZmNCED4*/ASI locus illustrates environmental conditionality with striking quantitative clarity: contributing-QTL PVE increases from 3.2% under well-watered conditions to 16.8% under severe drought as ABA-mediated pollen viability becomes the primary rate-limiting step in grain set under osmotic stress [20, 30], with direct implications for genomic-selection training environments, since models calibrated exclusively under optimal conditions may systematically underweight stress-responsive loci that are critical for performance in farmers’ marginal environments [9, 40, 89].

Haplotype phase, allelic architecture, and breeding strategy The practical breeding value of the identified hotspot loci depends critically on the allelic-phase configuration at linked loci within each hotspot [90–91]. Favorable alleles in coupling phase (on the same chromosome) are immediately deployable through standard marker-assisted backcrossing, whereas those in repulsion phase (on opposite chromosomes) require large segregating populations and multiple cycles of recombinant selection to recover favorable multi-locus genotypes at practical frequencies. The starch-biosynthesis cluster on chromosome 9 exemplifies the breeding advantage conferred by natural coupling-phase enrichment: long-term industrial selection for waxy starch simultaneously captured favorable alleles at waxy1, ae1, and su1 [65–66], enabling comparatively rapid and efficient variety development without deliberate pyramiding. Where linked resistance loci occur in coupling phase, concurrent resistance deployment is facilitated without the prohibitive population-size requirements imposed by repulsion-phase configurations [86–87]; determining the phase configuration at each resistance cluster in the target germplasm is therefore a prerequisite for efficient pyramiding and is a priority for follow-up haplotype analysis. For genomic regions in which favorable alleles may occur in repulsion phase (a configuration that, if confirmed by haplotype analysis, would apply to hubs such as the GIF1 grain-filling region on chromosome 5), breeders can employ several strategies: systematic screening of diverse germplasm panels and exotic collections for rare coupling haplotypes; designing large F or backcross populations to recover favorable recombinants; implementing haplotype-resolved genomic selection that explicitly models multi-locus haplotype effects rather than decomposing them into independent single-marker additive values; or applying precision genome editing for phase conversion at tightly linked loci where recombination-based decoupling is impractical [84–85]. Future integration of meta-QTL positions with high-density haplotype data from diverse germplasm panels could systematically classify all 23 hotspots by coupling-phase frequency, enabling rational, evidence-based stratification of deployment priorities from immediately actionable hotspots to longer-term investments requiring recombinant selection or genome-editing approaches [90–91].

Incorporating emerging pan-genome resources that capture presence–absence and structural variation, combined with long-read sequencing data capable of resolving complex genomic regions, could identify variants at hotspot loci that are currently invisible to SNP-based mapping [47].

### Cross-cereal conservation and translational opportunities

The recovery of confirmed orthologs for prioritized candidates across the comparison cereals (strongest for conserved developmental and metabolic genes) identifies substantial translational opportunities for the broader cereal research community [47]. The near-identity of Wx1/GBSSI across cereals (85–88% amino-acid identity) has already enabled waxy-starch programs in maize [92], rice [93–94], barley [95], and wheat [96] using analogous CRISPR gene-editing strategies that directly leverage functional knowledge gained in one cereal system to accelerate progress in others, demonstrating the directness and speed of the translational pathway when functional orthologs are well characterized. The ZmMATE2–SbMATE ortholog pair (∼84% amino-acid identity) [75, 97] enables direct syntenic marker transfer between maize aluminum-tolerance and sorghum acid-soil-adaptation breeding programs, with direct relevance to food security on aluminum-toxic Oxisols covering approximately 40–50% of arable tropical land in Africa and South America, where both maize and sorghum are grown extensively. The eIF4E broad-spectrum virus-resistance mechanism (>90% identity to barley rym4/rym5) represents the most directly translatable resistance mechanism identified in this study, with established functional characterization and breeding deployment in barley providing a detailed roadmap for analogous maize programs [98]. Conversely, the seven hotspot regions harboring candidates that lack detected orthologs in the comparison cereals under current database coverage whether owing to genuine evolutionary divergence or annotation incompleteness represent priority targets for synteny-guided ortholog mining and de novo functional characterization. These contrasting examples collectively position the cross-cereal ortholog framework as a powerful bidirectional knowledge-transfer resource.

## Conclusions

This meta-QTL atlas synthesizes three decades of maize quantitative genetics into a validated foundation that supports an actionable framework for precision breeding. By revealing multi-trait architecture, environmental context, and functionally coherent candidate genes, this study may enable geneticists and breeders to make more informed strategic decisions including selecting complementary loci for multi-trait improvement, deploying environment-specific alleles for targeted adaptation, managing antagonistic regions that create undesirable trade-offs, and prioritizing genome-engineering targets with maximum breeding utility. The framework offers a transferable template for similar efforts in other cereals, with pan-cereal conservation enabling reciprocal knowledge transfer across global crop-improvement programs addressing food security and climate adaptation.

## Declarations

### Ethics approval and consent to participate

Not applicable. This study is a meta-analysis of previously published quantitative trait locus (QTL) and genome-wide association study (GWAS) data; it did not involve any new experiments on humans, animals, or plants.

### Consent for publication

Not applicable.

### Availability of data and materials

All data generated or analyzed during this study are included in this published article and its additional files. The complete compiled QTL dataset (2,701 QTLs), the catalog of 187 meta-QTLs, hotspot composition, candidate-gene validation evidence, and cross-cereal ortholog tables are provided as Additional file 1 (sheets S1–S6). Source publications underlying the compiled dataset are cited within the compiled QTL dataset (Additional file 1) and in the reference list. No custom code was required beyond the publicly available software packages cited in the Methods (BioMercator V4.2.3; R).

### Competing interests

S.P. is employed by Corteva Agriscience. The authors declare that the research was conducted in the absence of any commercial or financial relationships that could be construed as a potential competing interest. T.R. and K.K. declare no competing interests.

### Funding

This research received no specific grant from any funding agency in the public, commercial, or not-for-profit sectors.

### Authors’ contributions

S.P. conceived and designed the study, performed the literature mining, meta-QTL analysis, candidate-gene prioritization, and cross-cereal ortholog analysis, and drafted the manuscript.

T.R. supervised the project, contributed to study design and interpretation, and critically revised the manuscript. K.K. contributed to interpretations and critically revised the manuscript. All authors read and approved the final manuscript.

## Acknowledgements

The authors thank the maize genetics community whose published QTL and GWAS studies made this meta-analysis possible, and the MaizeGDB, Gramene, and BioMercator development teams for the public resources used in this work.

## Abbreviations

ABA: abscisic acid
AIC: Akaike information criterion
ASI: anthesis–silking interval
BC: backcross
BLAST: Basic Local Alignment Search Tool
CI: confidence interval
cM: centiMorgan
CO: Crop Ontology
CRISPR: clustered regularly interspaced short palindromic repeats
DH: doubled haploid
FDR: false discovery rate
*GBSS*: granule-bound starch synthase
GWAS: genome-wide association study
LOD: logarithm of the odds
Mb: megabase pairs
MQTL: meta-quantitative trait locus
PVE: phenotypic variance explained
QTL: quantitative trait locus
RIL: recombinant inbred line
SNP: single-nucleotide polymorphism

## Figures

Each main figure is supplied as a separate high-resolution file (PDF, 300–600 dpi) per BMC Plant Biology requirements. Legends are listed below.

## Supplementary figure legends

Supplementary figures are provided as Additional files (TIFF). Legends are listed here for reference.

**Figure S1.** Chromosome-level distribution of meta-QTLs. Number of high-confidence meta-QTLs per maize chromosome (B73 RefGen_v5), showing how the 187 meta-QTLs are partitioned across the ten chromosomes and highlighting the chromosomes carrying the highest meta-QTL counts. MQTL, meta-quantitative trait locus.

**Figure S2.** Genome-wide QTL density across 10 Mb windows. Ideogram representation of the ten maize chromosomes with QTL density displayed as a heatmap across 10 Mb sliding windows; centromeres are marked. Chromosome 1 harbors the most QTLs, while chromosomes 9 and 10 exhibit the highest density per 10 Mb, and approximately 38.2% of QTLs localize to pericentromeric regions (±20 Mb from centromeres), reflecting suppressed recombination and extended haplotype blocks.

**Figure S3.** Shift in yield–stress co-localization frequency from optimal to drought conditions. Trait-class co-localization matrices among the five meta-QTL categories under optimal, drought, and low-nitrogen conditions. The proportion of yield (GYC) meta-QTLs overlapping physiology and stress-response (PHS) loci rises under drought, peaking at 71.4%, while GYC × PDA and GYC × GQN overlaps dominate under optimal conditions. GYC, grain yield and components; PDA, plant development and architecture; PHS, plant physiology and stress adaptation; GQN, grain quality and nutritional composition; DPR, disease and pest resistance.

**Figure S4.** Meta-QTL evidence and statistical support. Summary of the evidence layers underpinning the 187 high-confidence meta-QTLs, including the distribution of contributing QTLs per meta-QTL, GWAS co-localization support, dual-platform verification, and bootstrap positional stability, documenting the statistical robustness of the meta-QTL catalog. MQTL, meta-quantitative trait locus; GWAS, genome-wide association study.

## Additional files

Additional file 1. Supplementary tables (.xlsx). A single multi-sheet workbook containing all six supplementary data tables (sheets S1–S6); cited in text as “Additional file 1 (sheet S1)”, etc.

Sheet S1 (S1_All_QTLs): all compiled QTLs forming the raw input dataset (2,701 QTLs from 304 studies), with trait, source study, population, environment, chromosome, physical coordinates, LOD, PVE, R², confidence interval, and candidate-gene annotations.

Sheet S2 (S2_MQTL_Master): Master catalog of all 187 high-confidence meta-QTLs (B73 RefGen_v5), giving genetic position, 95% CI, physical interval, bin, summit and assembly, together with key traits, stress type, contributing-QTL and population counts, mean PVE, mean LOD, candidate genes, and a data-quality flag; this sheet also serves as the definitive per-MQTL coordinate reference.

Sheet S3 (S3_GWAS_Colocalization): Per-MQTL GWAS co-localization detail including coordinates, key traits, and the GWAS evidence supporting each co-localization.

Sheet S4 (S4_Hotspot_Detail): Composition of the 23 genomic hotspots, including supporting MQTLs, key orthologs/synteny, prioritized candidate genes (top-priority maize candidates with cross-cereal ortholog identity; ‘absent’ marks candidates lacking a detected ortholog in the comparison cereals), and the classification columns (Class, substantive trait-category count, dominant-pathway PVE, and maximum member-MQTL PVE).

Sheet S5 (S5_Gene_Validation): Validation evidence for the 119 characterized candidate genes: validation method, key references, evidence, breeding deployment, and confidence level.

Sheet S6 (S6_Ortholog_Full): Full cross-cereal ortholog table with BLASTp % amino-acid identity and references for rice, sorghum, wheat, and barley.

Additional file 2. Figure S1 (PDF). Chromosome-level distribution of meta-QTLs. Additional file 3. Figure S2 (PDF). Genome-wide QTL density across 10 Mb windows.

Additional file 4. Figure S3 (PDF). Shift in yield–stress co-localization frequency from optimal to drought conditions.

Additional file 5. Figure S4 (PDF). Meta-QTL evidence and statistical support.

Additional file 1 is a single multi-sheet workbook; its six sheets (S1–S6) are cited individually in the text as “Additional file 1”.

## Notes

### Competing Interest Statement

The authors have declared no competing interest.

### Summary of Updates

Revised analysis that includes figures and tables

