## Supplementary figures and images for "Multi-Trait Meta-QTL Analysis Reveals Genomic Hotspot Classes for Strategic Maize Improvement"

### Figure S1

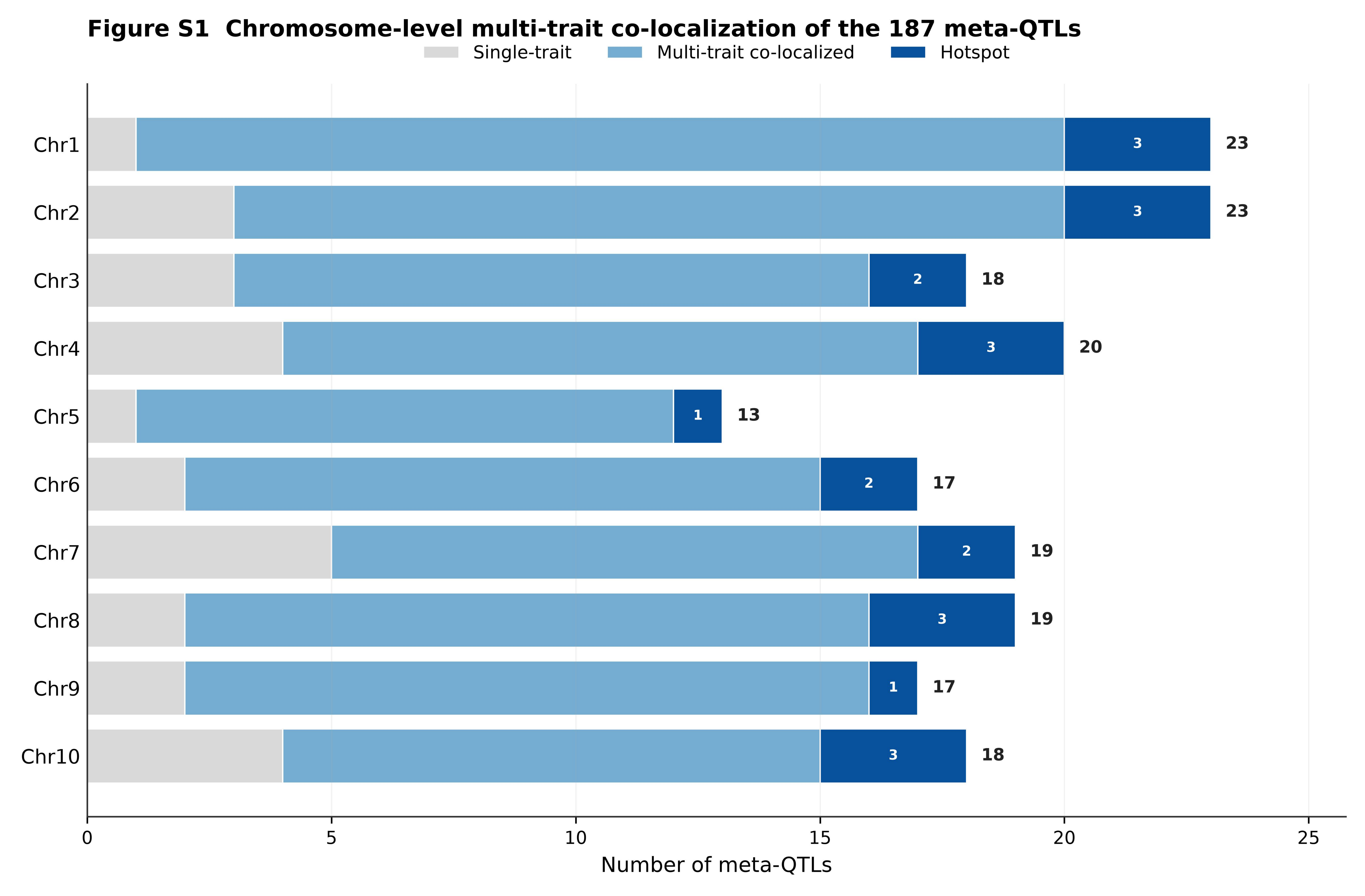

### Figure S2

#### Figure S2 Genome-wide distribution of QTL density in 10 Mb windows

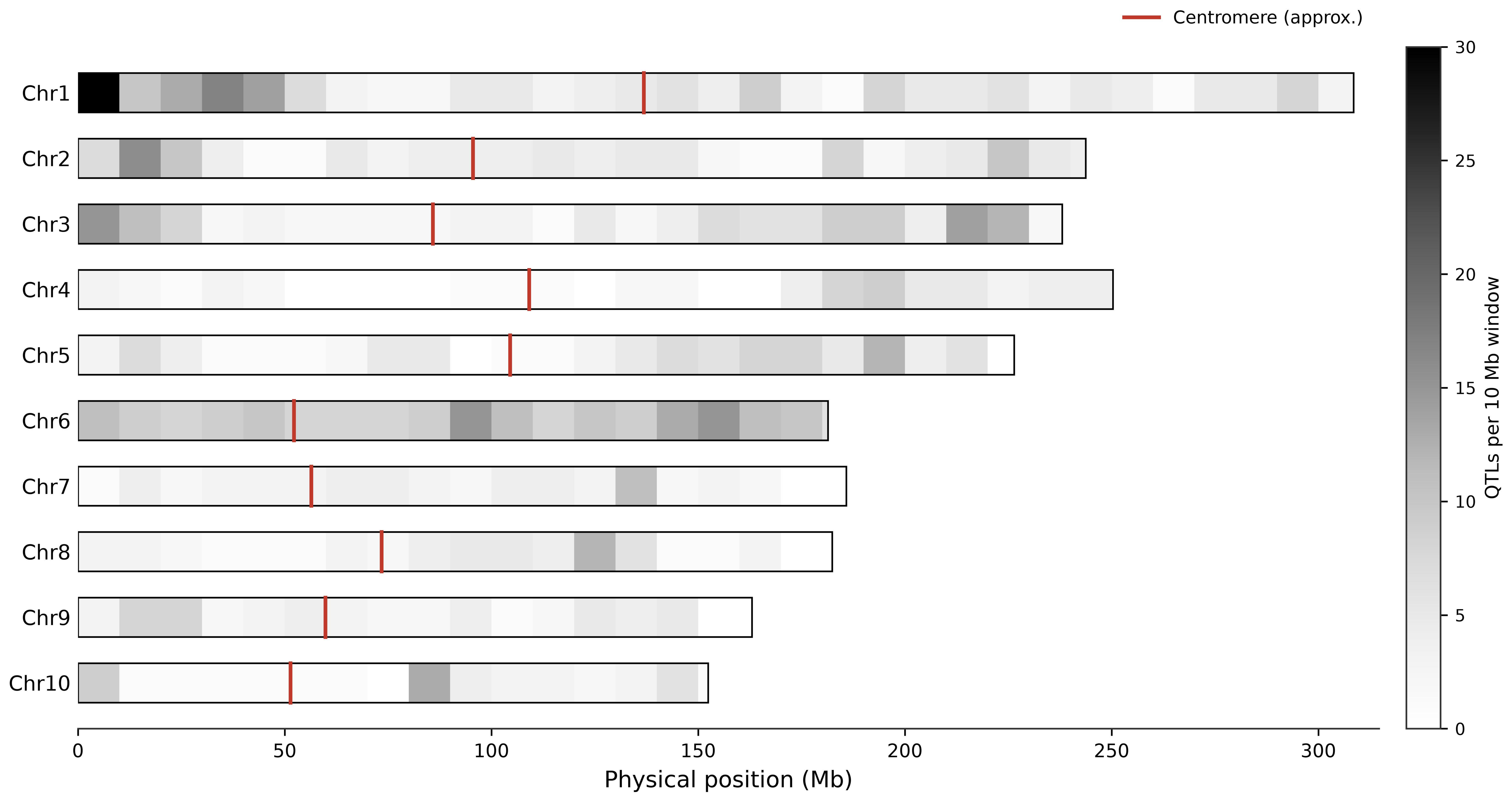

### Figure S4

**Figure S4 Evidence and statistical support of the 187 meta-QTLs**

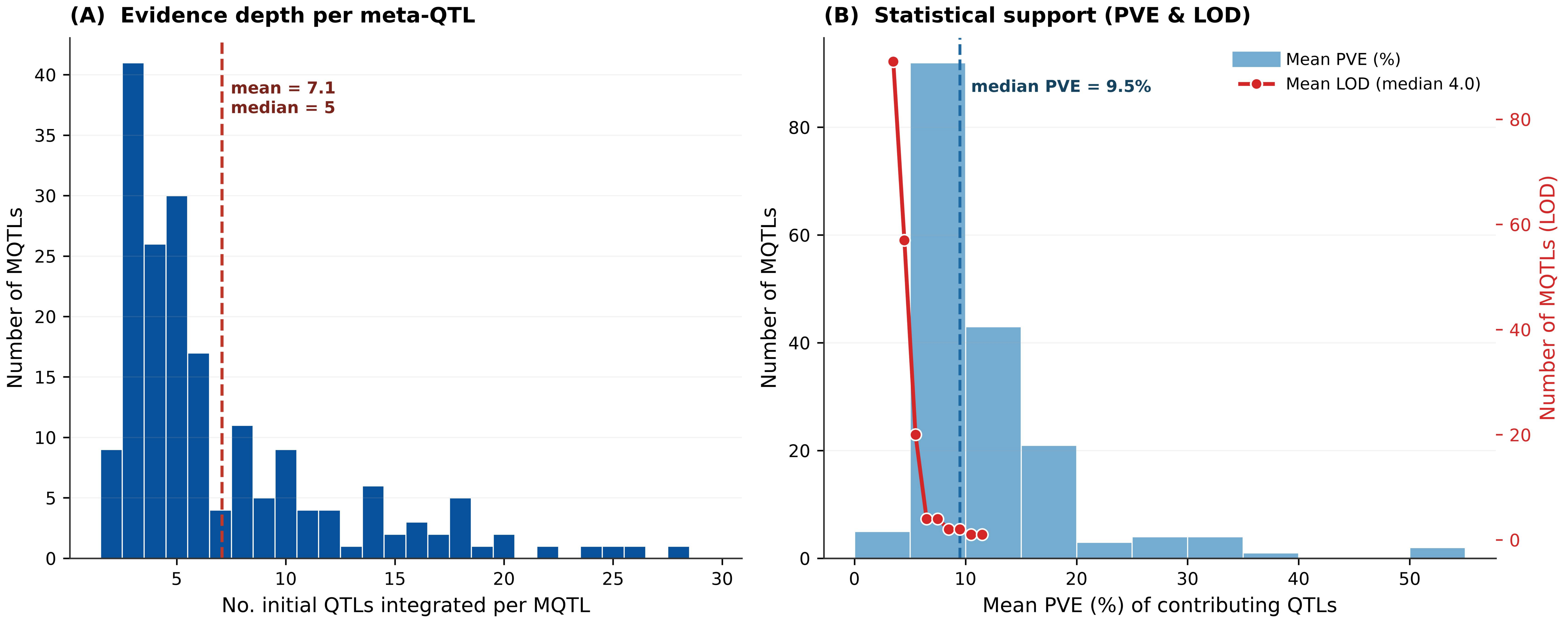
