## Supplementary material for "Multi-Trait Meta-QTL Analysis Reveals Genomic Hotspot Classes for Strategic Maize Improvement": Figure S3

**Figure S3  Yield-stress co-localization frequency from optimal to drought conditions**

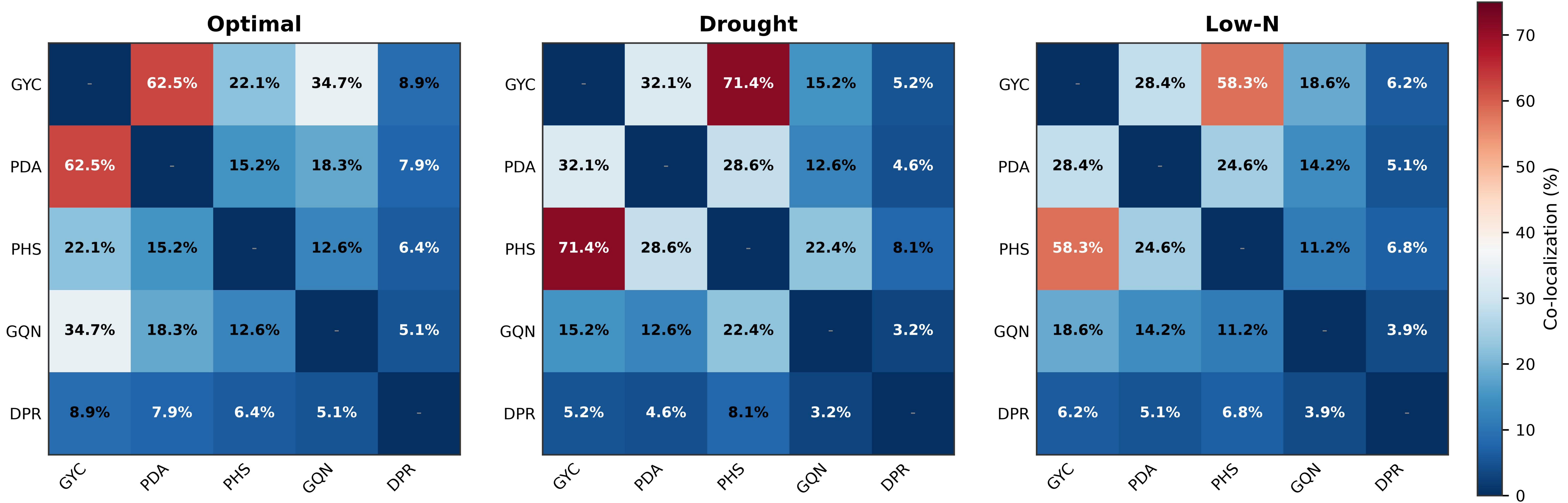


*GYC × PHS co-localization increases from 22.1% (optimal) to 71.4% (drought), revealing environment-specific genetic architecture.  GYC, grain yield & components; PDA, plant development & architecture; PHS, plant physiology & stress adaptation; GQN, grain quality & nutritional composition; DPR, disease & pest resistance.*
